# Large-scale changes of molecular network states explain complex traits

**DOI:** 10.1101/2022.12.05.519111

**Authors:** Matthias Weith, Jan Großbach, Mathieu Clement-Ziza, Ludovic Gillet, María Rodríguez-López, Samuel Marguerat, Christopher T. Workman, Paola Picotti, Jürg Bähler, Ruedi Aebersold, Andreas Beyer

**Affiliations:** Excellence Cluster on Cellular Stress Responses in Aging Associated Diseases, University of Cologne, Germany; Lesaffre Institute for Science and Technology, Lesaffre, Marcq-en-Baroeul, France; Department of Biology, Institute of Molecular Systems Biology, ETH Zürich, Switzerland; Department of Biotechnology and Biomedicine, Technical University of Denmark, Lyngby, Denmark; University College London, Institute of Healthy Ageing and Department of Genetics, Evolution & Environment, London, United Kingdom

## Abstract

The complexity of many cellular and organismal traits results from poorly understood mechanisms integrating genetic and environmental factors *via* molecular networks. Here, we show when and how genetic perturbations lead to molecular changes that are confined to small parts of a network *versus* when they lead to large-scale adaptations of global network states. Integrating multi-omics profiling of genetically heterogeneous budding and fission yeast strains with an array of cellular traits identified a central state transition of the yeast molecular network that is related to PKA and TOR (PT) signaling. Genetic variants affecting this PT state globally shifted the molecular network along a single-dimensional axis, thereby modulating processes including energy- and amino acid metabolism, transcription, translation, cell cycle control and cellular stress response. We propose that genetic effects can propagate through large parts of molecular networks because of the functional requirement to centrally coordinate the activity of fundamental cellular processes.

**Graphical abstract:** Graphical abstract:
Genetic variants directly or indirectly affecting the activity of PKA and/or TOR signaling cause global changes of transcriptomic and proteomic network states by modulating the activity of diverse cellular functions and network modules. Using marker genes acting downstream of PKA and TOR signaling we are able to quantify the activity status of combined PKA and TOR signaling (‘PT Score’). This PT Score correlates with major transcriptomic and proteomic changes in response to genetic variability. Those large-scale molecular adaptations correlate with and explain phenotypic consequences for multiple cellular traits. Variants affecting the stoichiometry of proteins within a specific module have regional effects that remain confined to smaller parts of the molecular network. Variants affecting only one or very few proteins change molecular networks only locally. The global reorganization of network states caused by variants of the first type result in consequences for many cellular traits (i.e. pleiotropic effects), such as growth on different carbon sources, stress response, energy metabolism and replicative lifespan. (Created with BioRender.com)

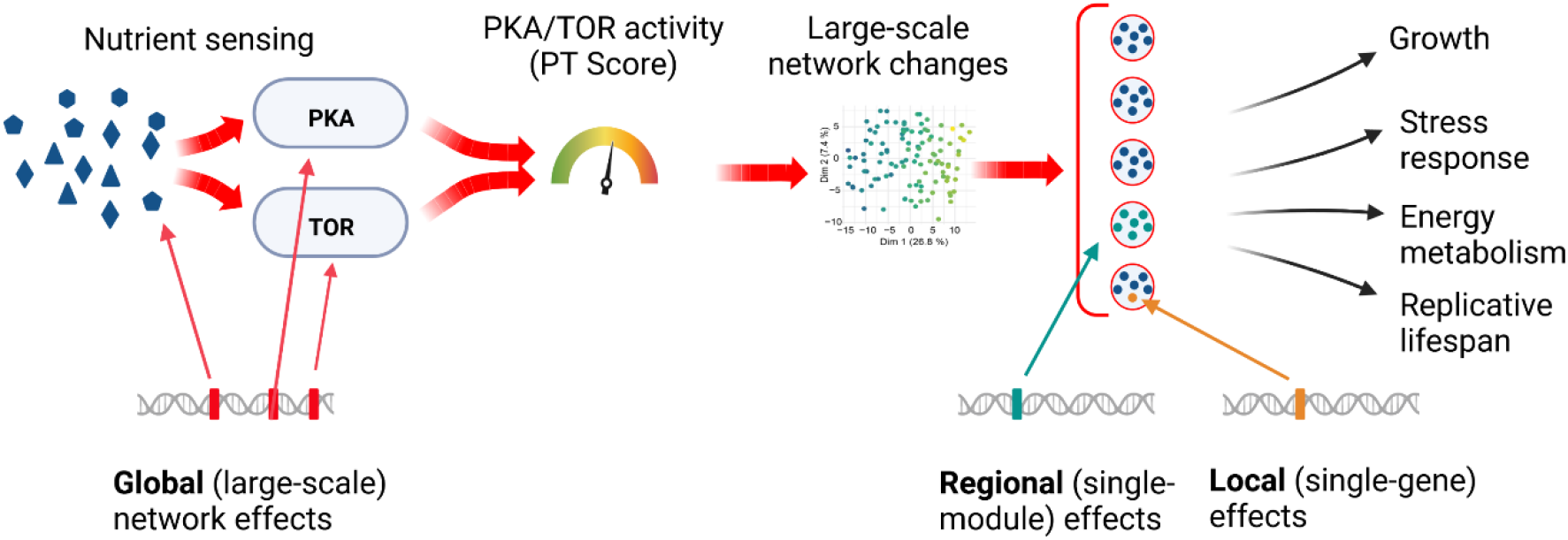

## Introduction

The genetic complexity of traits such as human body size and disease susceptibility has been well known for many years. Still there is a lack of understanding about how the ensemble of trait-associated genetic variants and environmental factors are integrated at the molecular level. Recent work has proposed an “omnigenic” as opposed to a “polygenic” model for the genetic architecture of complex traits in humans, in principle stating that modulation of any gene that is expressed will have non-zero effects on all traits associated with a given tissue (Boyle et al., 2017). Central to the omnigenic model is the notion that genetic variants can affect “peripheral genes” and, via mediation by the cellular gene-regulatory network, ultimately affect trait-determining “core genes” (Liu et al., 2019). However, the mechanistic basis for the transmission of such effects across the network has remained unclear. In contrast, work focusing on the modular sub-structure of molecular networks favors the view that specific sub-networks or network modules are associated with disease phenotypes (Chen et al., 2008; Han, 2008; Peters et al., 2017; Schadt, 2009; Vidal et al., 2011). Fundamental to the modularity of a molecular network is the notion that cellular functions require particular stoichiometries between molecular components. This concept facilitates mechanistic understanding of the propagation of genetic variant effects since members of a network module are often subject to co-regulation. In this latter framework however, stoichiometry and co-regulation appear largely confined to the boundaries of individual network modules and consequently, variant effects would have limited reach. So far, no concept has been established that includes coordination across separate functional modules as the basis for genetic variant effects.

Here we propose a model that explains complex and far-reaching genetic effects as alterations of cellular network states. We define a network state as a particular molecular configuration of a cell, i.e. its specific composition of transcripts, proteins, other (small) molecules and their molecular states as defined by e.g. protein post-translational modifications. By “network” we refer to the entire ensemble of interactions occurring between these components within a cell. Our model is based on the assumption that only a small fraction of all theoretically possible network states are physiologically feasible and favorable. This subset of network states is defined by natural constraints, which may reflect basic biophysical principles, such as the need to balance anabolic and catabolic activities to preserve homeostasis, cellular limits such as maximum proteome size, molecular crowding or the availability of membrane space as well as parameters such as protein cost (Frei et al., 2020; Hu et al., 2020; Kleijn et al., 2022; Molenaar et al., 2009; Mori et al., 2021). Yet other constraints evolve by adaptation to the ecological niche (Balakrishnan et al., 2021). Accordingly, also in the context of evolution network states are constrained to remain within a relatively small sub-space of all possible states, exemplified by the fact that most genomic positions in coding regions are under purifying selection (Ellegren, 2005). Identification and quantification of natural constraints requires systematic and quantitative description of global network state adaptations to different conditions, as has recently been demonstrated for *E. coli* (Mori *et al*., 2021). In this organism, seminal work also elucidated mechanisms of proteome allocation, providing an intriguing example for constrained cellular network behaviour (Basan et al., 2015; You et al., 2013).

Network states can change through sensing the cell’s environment or internal molecular state. Control mechanisms exemplified by signaling pathways ensure that the network remains in a viable state while adjusting a multiplicity of cellular functions to a particular environment. Genetic variants can shift network states within the boundaries of viability and can act on network state-modifying mechanisms in different ways: by affecting sensing or signaling, but also by modifying the state of (intermediate) metabolites being sensed. Using this concept as the basis for our model, we propose a new classification of genetic effects: (1) variants affecting only a very small part of a network, such as a single protein or protein complex (subsequently called ‘local effects’), (2) variants affecting single or grouped modules of the network, such as individual signaling, regulatory or metabolic pathways (subsequently called ‘regional effects’) and (3) variants affecting the balance between seemingly unrelated network modules. The latter variants change particular aspects of the molecular configuration of a cell, which require a far-reaching adjustment of the cellular metabolism and module activities (subsequently called ‘global effects’). Here, we present examples of all three types of genetic variants.

We set out by applying this paradigm to a complex cellular phenotype, namely the ability of yeast to efficiently overcome temporary temperature stress. Heat stress is among the best studied perturbations in yeast (Gasch et al., 2000). Many aspects, such as organization of the proteome into liquid phases (Wallace et al., 2015) are conserved in human cells (Franzmann and Alberti, 2019). Hence, it remains of great interest to understand how genetic variation influences thermotolerance and cellular stress resistance in general. Using a collection of yeast segregants, we studied the genetic contribution to efficient outgrowth after temporary heat stress, which we refer to as ‘heat resilience’. Transcriptomic, proteomic and phosphoproteomic measurements were employed to comprehensively chart the molecular network state in each segregant strain (Grossbach et al., 2022). We found genetic variants with effects on specific, heat-stress related proteins and others that determine resilience through a broad cellular program that is closely related to PKA and TOR signaling. The network state that results from these signaling activities (PKA/TOR-related or PT network state) can be summarized as a single quantitative trait. We characterize the global quantitative alterations of transcripts, proteins, phosphorylation and metabolic features that constitute changes of this network state across a wide range of environmental and genetic influences. Finally, we demonstrate the conservation and phenotypic relevance of the PT network state across large evolutionary distances in other yeast species and in the presence of oxidative stress. Due to the utmost importance of cellular processes under the control of Ras and TOR signaling, we envision that similar global network changes underly variability of complex human traits including health span and disease susceptibility.

## Results

### 1. Genetic and molecular mapping of heat resilience

Previous studies of thermotolerance in yeast found genetic determinants that convey advantages for growth under persisting high temperature (Abrams et al., 2022; Caspeta et al., 2016; Sinha et al., 2008; Sinha et al., 2006; Steinmetz et al., 2002; Weiss et al., 2018; Yang et al., 2013) and for heat shock survival (Gibney et al., 2013; Jarolim et al., 2013). Here, we aimed to understand which factors facilitate the recovery of yeast following a short, sub-lethal heat stress episode. We expected that a range of dynamic processes during and following transient heat stress collectively determine the time required to re-establish maximum growth rates, the ‘heat-induced lag’. Since the extent of damage and the initial response to the heat insult will be determined by the molecular configuration of the cell during unperturbed growth, this setup probes a different aspect of thermotolerance as compared to persisting high temperature and heat shock survival. To find genetic determinants of the thermotolerance trait, we made use of a well-studied cross between isogenic haploid derivatives of the common lab strain S288c, BY4716, and the vineyard-isolate strain RM11-1a (BYxRM collection (Brem et al., 2002)). We exposed exponentially growing cultures of the parental strains and of 100 segregants to transient heat stress at 45 °C for 8 min, which we confirmed to be sub-lethal (Fig. S1A). Following heat stress or mock treatment, samples of the cultures were diluted into fresh medium and growth curves were recorded (Fig. 1A). From these curves, strain-specific growth characteristics were inferred and used to estimate the heat-induced lag (Fig. 1B).

**Figure 1:**
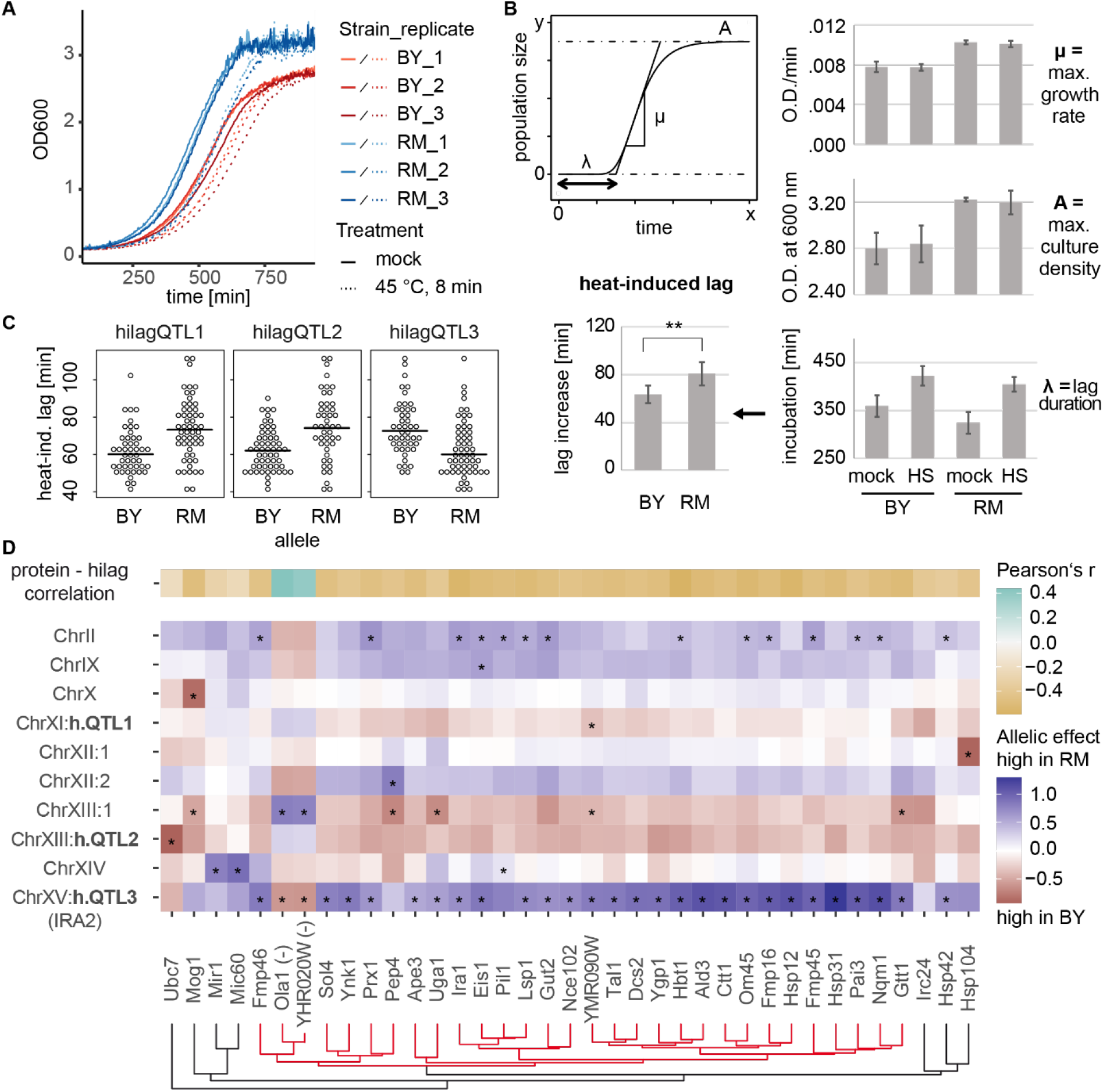
Identification of genetic and molecular determinants of heat-induced lag. **A** Growth curves recorded for BY4716 and RM11-1a after transient heat stress treatment (45 °C, 8 min) and dilution into fresh growth medium. **B** Quantification of growth parameters. Heat-induced lag is calculated as the difference in lag duration (λ) between heat-treated (HS, 45 °C, 8 min) and mock-treated samples for each strain. **C** Distribution of hilag measurements for strains carrying opposing parental alleles at significant loci identified by QTL mapping (15 % FDR, see Fig. S1C). **D** Mapping of heat-induced lag to protein abundances and overview of corresponding pQTL. 37 proteins passing a 20 % FDR threshold as predictors of heat-induced lag are shown. The top row shows correlation of the protein’s abundance with heat-induced lag. Fields in all other rows show allelic effects (difference in mean protein abundance between strains carrying the RM allele compared to the BY allele at the indicated loci) and asterisks indicate significant pQTL-target relations as described in (Grossbach *et al*., 2022). Columns or proteins were ordered according to hierarchical clustering. Abundance scaling of Ola1p and YHR020W (marked by “(-)”) was inverted across the segregants for clustering. h.QTL = hilagQTL.

The parental strains scored close to the median (66 min) of all measurements of heat-induced lag across the collection (BY: 64 min, RM: 81 min, Fig. S1B), consistent with a transgressive pattern of segregation and presumably, a polygenic basis. To localize genetic factors that contribute to the variation of heat-induced lag, we performed mapping of quantitative trait loci (hilagQTL) using a Random Forest-based approach (Michaelson et al., 2010). Three loci passed a threshold of 15 % FDR (Fig. S1C) and showed variable effect directions (Fig. 1C), partially accounting for transgressive segregation. Each locus explained a difference of about 10 min in the duration of heat-induced lag. An additive random-effects model based on the most significant markers at these loci explained up to 35 % (adjusted R^2) of trait variability across the strains. Based on the parental replicate measurements, we estimated the heritability of heat-induced lag to be 46 % (adj. R^2^), leaving about a quarter of the estimated heritability unexplained by effects at these QTL. Regarding potential mediators of the QTL effects, we noted that the locus on Chromosome XV (hilagQTL3) contained *IRA2*, which is known to induce widespread changes of transcriptional and cellular traits in this yeast cross (Nguyen Ba et al., 2022; Smith and Kruglyak, 2008). Replacing *IRA2* by the RM allele in the BY background indeed approximated the effect of the QTL (Fig. S1D).

To identify proteins which might have affected heat-induced lag duration, we compared protein abundances recorded for each strain (Grossbach *et al*., 2022) to trait differences across the collection. Specifically, we trained a Random Forest model predicting heat-induced lag as a function of protein abundance. Setting a permutation-based false-discovery rate (FDR) threshold of 20 %, we identified 37 predictive proteins (Fig. 1D). This protein set contained candidates that are well known contributors to heat shock survival, such as Hsp104 (Cherkasov et al., 2013) and Ctt1 (Davidson et al., 1996). We further asked which genetic loci would be associated with variation in predictors of heat-induced lag by assigning protein abundance QTL (pQTL) to this set of proteins. This approach revealed loci with strong effects on few, individual proteins. For example, a pQTL on Chromosome XII affected levels of Hsp104, presumably in *cis*, since it was found close to the *HSP104* locus. Mog1, a nucleotide-release factor for the Ran GTPase Gsp1 that participates in the osmotic stress response (Lu et al., 2004), was the sole significant target of a pQTL on Chromosome X. On the other hand, a subset of 30 out of the 37 hilag predictor proteins exhibited highly correlated changes across the strain collection (red cluster in Fig 1D), 28 of which shared a significant pQTL at the *IRA2* locus (hilagQTL3). We further noticed that genetic loci often had opposing allelic effects across many of these proteins: whereas the RM alleles at the loci ChrII, ChrIX, ChrXII:2, and hilagQTL3 elevated levels of these proteins, the RM alleles at the loci hilagQTL1, ChrXIII:1 and hilagQTL2 reduced levels of the same proteins. These effects were consistent with the effect directions of the three hilagQTL on heat-induced lag (Fig. 1C). Hence, these 30 proteins seemed to be under the control of a common regulatory program. Several genetic variants affected this putative regulatory program, resulting in a convergent effect on heat-induced lag.

Taken together, our mapping results suggested that differences in the extent of heat-induced lag and its transgressive segregation in this cross can be explained both by variants that change the abundance of individual proteins such as Hsp104, as well as by variants that affect a more general program of gene expression. In particular, mapping the trait to protein abundances indicated that the *IRA2* locus impacted a set of proteins with coordinated expression, which was additionally modulated by hilagQTL1, hilagQTL2 and potentially other loci.

### 2. Quantification of a PKA/TOR-related program of gene expression

The *IRA2* locus (coinciding with hilagQTL3) is a pQTL hotspot, i.e. a genomic region containing significantly more pQTL than expected by chance (Grossbach *et al*., 2022). Our analysis from above suggested that target proteins of this locus are subject to a common regulatory program. In order to corroborate this notion, we controlled for the effect of *IRA2* in the following way. First, we split the segregants into two groups depending on their *IRA2* alleles and analyzed these groups independently. Thus, we removed effects of the *IRA2* allele and partially removed effects of neighboring variants in linkage disequilibrium (LD). Next, we quantified the pairwise protein abundance correlation for all 225 *IRA2* target proteins (pQTL targets at 10 % FDR) within the two groups of strains. We speculated that, if *IRA2* target proteins were affected by additional loci, there should be a remaining correlation that is greater than random. This was indeed the case, as abundance-matched but otherwise random sets of proteins exhibited lower average pair-wise correlations (mean R^2^ = 0.06) than targets of the *IRA2* locus (mean R^2^ = 0.16, empirical p value < 1E-3 as determined by repeated sampling of random sets of proteins, Fig. S2A). We applied the same test for coordination among their targets to 11 other hotspots of protein abundance regulation (Fig. S2B). The degree of coordination among targets of the *IRA2* hotspot was larger than for 4 other hotspots with more than 100 targets (Fig. S2C). In sum, hotspots often affected network modules that remained coordinated across multiple genetic perturbations. However, the *IRA2*-related program encompassed an exceptionally large set of strongly coordinated proteins.

Ira2 is a GTPase-activating protein (GAP) that negatively regulates Ras1/2, which are upstream regulators of the PKA signaling pathway. The RM allele of *IRA2* is more active compared to the BY allele due to coding sequence differences (Nguyen Ba *et al*., 2022; Smith and Kruglyak, 2008). Hence, the effects of the *IRA2* locus on protein abundances likely result from genetic influence on PKA activity. The remaining coordination among *IRA2* targets in strains with identical alleles at this locus suggested that more loci could affect the same set of proteins. For example, a region on Chromosome XIII, which includes the *BUL2* gene, had strong and opposite effects across many *IRA2* targets (Figure 1D). Bul2 is involved in the endocytosis of amino acid permeases (Abe and Iida, 2003; Merhi and Andre, 2012) and thereby likely affects TORC1 signaling (here referred to as TOR signaling for simplicity) (Kwan et al., 2011). Thus, the same set of co-regulated target proteins may be affected by genetic variants altering the activities of PKA and TOR signaling. Strong crosstalk between the TOR and PKA pathways has been shown repeatedly (Chen and Powers, 2006; Ramachandran and Herman, 2011; Zhang et al., 2011), including signaling connections via mediators such as Mpk1 (Soulard et al., 2010) as well as convergence on common downstream effectors such as Rim15 (Lee et al., 2013; Pedruzzi et al., 2003; Reinders et al., 1998; Swinnen et al., 2006). PKA and TOR signaling also commonly affect a range of transcription factors including Gis1, Hsf1, Msn2/4, Sfp1, Dot6, Tod6 and others (Kunkel et al., 2019; Lee *et al*., 2013; Lippman and Broach, 2009). These observations led us to hypothesize that a large part of the protein abundance changes explaining the extent of heat-induced lag was under the common control of these two major pathways.

In order to corroborate the relevance of PKA and TOR signaling in this context, we compared our collection-wide protein abundance data to transcript abundance changes following chemical inhibition of the PKA and TOR pathways (Kunkel *et al*., 2019). Indeed, we observed that the effect of the RM allele of *IRA2* in our dataset was highly correlated to the effect of chemical PKA and TOR inhibition (Pearson’s r = 0.75 and 0.60, respectively, both significant at p < 1E-15, Fig. 2A). We next sought to quantify the status of PKA/TOR (PT) signaling in a given population of yeast cells. We therefore conceived a score that summarizes overlapping outcomes of PKA and TOR activity (‘PT score’) based on a set of 47 PT-induced and 44 PT-repressed genes selected by thresholding and filtering the chemical inhibition data. We then made use of the overlap between these markers and our proteomic data for the BYxRM cross to select 18 and 22 abundance markers for induced and repressed PKA/TOR activity, respectively (Fig. 2A). The PT score is the difference between the median abundance of proteins (scaled and centered across the collection) in the PT-induced set and the median of the PT-repressed set in a specific yeast strain. Thus, the PT score quantifies the activity of PKA/TOR signaling in a given strain relative to the collection-wide average.

**Figure 2:**
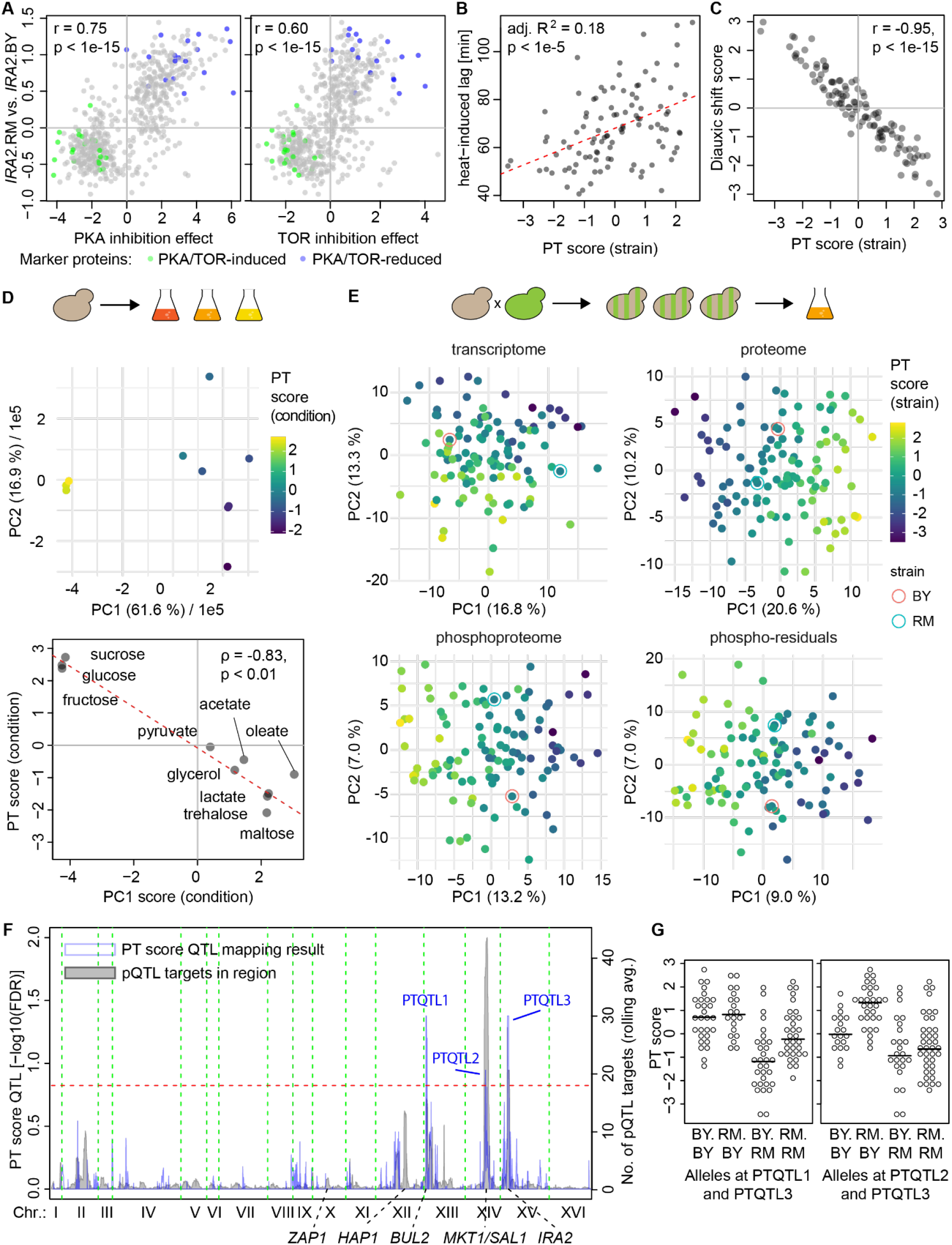
Scoring and genetic mapping of a PKA- and TOR-signaling-dependent network state. **A** Comparison of the effects of chemical inhibition of PKA (left) and TOR (right, both after treatment for 20 min, data from (Kunkel *et al*., 2019)) on transcripts to difference according to parental *IRA2* allele on the corresponding protein abundances in the BYxRM collection. Assignment of transcripts to PT-induced and -reduced sets is indicated by green and blue dots, respectively. **B** Correlation of PT score with heat-induced lag measurements per strain of the BYxRM collection. **C** Comparison of a score based on transcript level changes during the diauxic shift (Brauer *et al*., 2005) to the PT score for all strains of the BYxRM collection. **D** PCA of proteomic data for BY4742 grown on different carbon sources (Paulo *et al*., 2016) and comparison to the PT score for each sample, calculated based on protein abundances after scaling and centering across conditions, yielding a comparison to the global mean for the studied range of carbon sources. Upper panel: PC1 and PC2 of the PCA. Coloring by PT score. Fractions of total variance explained by each PC is indicated. Lower panel: Comparison of sample scores on PC1 to the PT score. Dashed red line shows the linear regression model (see main text). **E** PCA for molecular variability between strains of the BYxRM collection based on transcript, protein and phosphopeptide abundances as well as phospho-residual levels (see methods). Coloring by PT score. The PT score explained up to 86 %, 80 % and 68 % variability (adjusted R^2^) between segregant scores along PC1 of PCAs based on protein, phosphopeptide and phosphopeptide-residual abundances, respectively. For the transcriptome, the PT score correlated more strongly with the second PC (adj. R^2^ = 0.44) **F** QTL mapping of the PT score calculated for each segregant (blue curve) and overlay of the number of pQTL targets by marker (grey curve; rolling average of pQTL number across windows comprising 20 genetic markers). Dashed red line represents 15 % FDR threshold for QTL mapping. Known variants close to pQTL hotspots are indicated. **G** Distribution of PT scores for strains carrying indicated combinations of parental alleles at significant QTL for the PT score.

Consistent with the direction of effect of the *IRA2* allele (Fig. 1C), the PT score was positively correlated with heat-induced lag and explained up to 18 % variability (linear model adjusted R^2^, p < 1E-5, Fig. 2B), exceeding the effect of any single QTL identified in our RF mapping approach, including the *IRA2* locus itself. Further, the PT score and heat-induced lag still appeared to correlate after correcting for all three hilagQTL (partial correlation R’ = 0.19, p = 0.06) indicating that additional variability in the molecular network that determines heat-induced lag was captured by the PT score. As noted above, we observed inverse effects of the other hilagQTL on *IRA2* targets. Consistently, hilagQTL1 and hilagQTL2 had significant and inverse effects on the PT score (mean difference between RM and BY allele-carrying strains: +0.59 and +0.67, respectively, both p < 0.05) as compared to hilagQTL3 (−1.19, p < 1E-5).

### 3. A network state model to explain far-reaching genetic effects

The observations above indicated that several genetic variants jointly modulated a coordinated program of gene expression that resembled the effects of PKA and TOR signaling. We have previously shown that a pyruvate kinase variant modulated the metabolic balance between fermentation and respiration during exponential growth in fission yeast (Kamrad et al., 2020). Given the roles of PKA and TOR in regulation of the transition between these metabolic states at the diauxic shift (Pedruzzi *et al*., 2003), we hypothesized that the PT score might quantify a related balance. Hence, for comparison, we extracted a set of markers based on transcript abundance changes during the diauxic shift (Brauer et al., 2005), analogous to the markers for the PT score (31 diauxic shift-induced and 30 -reduced proteins). The difference in medians between these marker sets showed almost perfect correlation with the PT score across BYxRM segregants (Fig. 2C, r = −0.95, p < 1E-15). This correlation indicated that quantifying the outcome of genetic effects in this collection in terms of the PT score was similar to placing individual strains on a ‘diauxic shift-axis’. Hence, the gene expression program represented in the PT score resembled the program initiated at the diauxic shift, i.e., transiting from fermentation to respiration.

It is known that adaptation of large parts of the molecular configuration of yeast cells due to changes in nutrient source involves the PKA and TOR signaling pathways (Broach, 2012). We therefore asked whether our quantification of the state of these two pathways would accurately capture nutrient-dependent changes. To test this, we analyzed a proteomic dataset generated for the S288c-derived strain BY4742 grown on 10 different carbon sources (Paulo et al., 2016). We calculated PT scores for each condition based on the abundances of 40 and 39 proteins that overlapped with the PT-induced and -reduced marker sets, respectively, as derived from chemical inhibition data (see Methods). The highest scores were attributed to populations grown on glucose, sucrose and fructose, as expected (Paulo *et al*., 2016), while other carbon sources led to lower PT scores for the respective populations. We performed principal component analysis (PCA) to reduce the complexity of the proteomic changes to a low-dimensional space (Fig. 2D). We then compared PT scores calculated for the individual growth conditions to their respective placement along the first principal component (PC1). These two quantifications of the proteomic state indeed showed strong correlation (ρ = −0.83, p < 0.01, Fig. 2E). Only the sample grown on oleate deviated noticeably from the global trend. After exclusion of this outlier, the predictive accuracy of the PT score for PC1 scores in a linear model across conditions reached up to 96 % (adj. R^2^, p< 1E-5). Since PC1 scores in this dataset reflected more than 60 % of total proteome variability, we concluded that for a broad range of carbon sources, PT score differences correctly captured the major adaptation of the proteome.

Consequently, we asked whether the PT score was predictive for larger parts of proteome variation caused by genetic variation. Indeed, the PT score correlated with a major fraction of overall proteome variation in the BYxRM collection. For example, the PT score strongly correlated with the first dimension in a PCA of the BYxRM proteomes (Fig. 2F).

Overall, linear models using the PT score reached proteome-wide significance (FDR < 0.05) for 622 of 1862 (33 %) protein abundances, explaining on average up to 22 % of their total variation (mean adj. R^2^ in a linear model). Including the *IRA2* and *BUL2* alleles simultaneously with the PT score in a combined correlation model only marginally reduced the number of significant partial PT score correlations to 508 (27 %, FDR < 0.05). Only 23 of these proteins were not significantly predicted by the PT score alone. Beyond protein abundance, the PT score was also predictive for substantial variation in transcriptome and phospho-proteome data from the same strain collection (Fig. 2F). Hence, the PT score captured major parts of molecular differences across the BYxRM collection in a single quantitative value. Conversely, the configuration of the cellular molecular network was shifted along an axis defined by this scalar. This seemed to be best explained by thinking of differences in PKA/TOR-related gene expression as a shift of the global network state.

### 4. Genetic mapping of the PT network state in the BYxRM collection

Next, we sought to identify genetic variants that affected the PT network state across the BYxRM collection. Strikingly, no segregant showed a PT score that fell between those of the parental strains (BY: −0.68, RM: −0.69), indicating highly transgressive segregation of this trait. We performed QTL mapping as above (Fig. S1C) using the PT score as a target trait and detected three regions containing predictive markers (PTQTL at 15% FDR, Fig. 2G). The RM allele at the most significant marker in PTQTL3 on Chromosome XV, which was close to *IRA2* (< 10 kb), was associated with a lower mean PT score (−1.41, p < 1E-8). This is consistent with the known role of Ira2 in Ras/PKA signaling and the stronger PKA-inhibitory effect of the RM variant (Smith and Kruglyak, 2008), suggesting that *IRA2* mediated the effect at this locus. PTQTL1 on Chromosome XII coincided with *BUL2*. The RM allele at the most significant marker was associated with a higher PT score (+0.58, p < 0.05). Strains carrying the RM allele at the most significant marker for PTQTL2 also had a higher mean PT score (+0.63, p < 0.05) than strains carrying the BY allele. This region contains known variants in *MKT1* and *SAL1*, which contribute to mitochondrial genome instability and consequently, a higher proportion of petite cells in populations of BY compared to RM (Dimitrov et al., 2009). There was considerable epistatic interaction between these PTQTL, which underlines the complexity of the PT network state (Fig. 2H). Specifically, the effect size of PTQTL1 (close to *BUL2*) was higher in strains carrying the RM allele at PTQTL3 (close to *IRA2*, +1.10) than in strains carrying the BY allele (+0.21, p < 0.05 for the interaction term), whereas the effect size at PTQTL2 (close to *MKT1/SAL1*) was higher in strains carrying the BY allele (+1.14) than in those carrying the RM allele (+0.24, p < 0.05 for the interaction term) at PTQTL3. Potentially related interactions between major pleiotropic effect loci in determining cellular fitness traits have been described recently (Nguyen Ba *et al*., 2022).

We observed that PTQTL signals often coincided with a high number of pQTL linking to the same region (Fig. 2G), which suggested to us that genetic influence on the PT network state might be more widespread across the genome. Hence, we evaluated the coordination between 40 PT marker proteins by computing allelic effects for all genetic markers in the BYxRM cross, irrespective of their significance (Fig. S2D). The PT score marker proteins fell into two groups, depending on whether they were positively or negatively affected by PKA/TOR inhibition (see above). Consistent with a general coordination between these proteins, we observed that alleles associated with increased abundance of one set of marker proteins had negative effects on the other (Fig. S2D). The largest differences were observed at markers coinciding or in linkage disequilibrium with hotspots of protein abundance regulation. Indeed, 7 of 12 pQTL hotspots (as detected in (Grossbach *et al*., 2022)) showed significant PT score differences at 5 % FDR and proteome-wide effects of pQTL hotspots were often correlated with the effects of chemical inhibition of PKA and TOR (Fig. S3A). On the other hand, 423 (of 2,078 in total) pQTL linkages originated at loci that caused very little change of the PT score (difference between marker set medians < 0.15). Among these loci with small associated differences in the PT score were the pQTL for Hsp104 and Mog1, which are protein predictors of heat-induced lag (Fig. 1F). In sum, these observations indicated that the PT network state was modulated by a wide variety of genetic variants, underlining the complexity of this trait. However, there were also a number of genetic variants not changing the PT state in a detectable way despite significantly affecting individual or groups of proteins.

### 5. Effect of PT network state differences on functional modules

The analyses above suggested that genetic variants which alter the PT network state have far-reaching effects, consistent with the coordination of a large diversity of cellular processes by the PKA/TOR-related regulatory program. In order to characterize these alterations in cellular processes and functions, we tested for correlation of the PT score with protein abundances and phosphorylation across the SGD-curated set of Gene Ontology (GO) slim terms (Cherry et al., 2012). The PT score was a significant linear predictor (q < 0.05) for the abundance of more than half of the assigned proteins in close to a third of all tested GO terms (29 of 95 terms with at least 3 members; Fig. 3A). Strong and highly directional correlations with the PT score comprised well-known targets of PKA and TOR signaling (Conrad et al., 2014), including cytoplasmic translation, ribosome biogenesis and related processes among positively correlated terms (Fig. 3A). Carbohydrate transport and to a lesser extent amino acid transport were also positively correlated with the PT score, consistent with the role of glucose transport capacity for glycolytic activity (Elbing et al., 2004). Further, we found many positive correlations between the PT score and proteins involved in transcription by RNA polymerase I and III but not II, in line with recent observations by others (Kleijn *et al*., 2022). Negative correlation was predominant within GO terms such as response to starvation and oxidative stress, cellular respiration, carbohydrate metabolism, generation of precursor metabolites and energy, and sporulation. Moreover, we found relatively high prediction accuracy but variable directionality of correlation with proteins annotated for protein phosphorylation and dephosphorylation.

**Figure 3:**
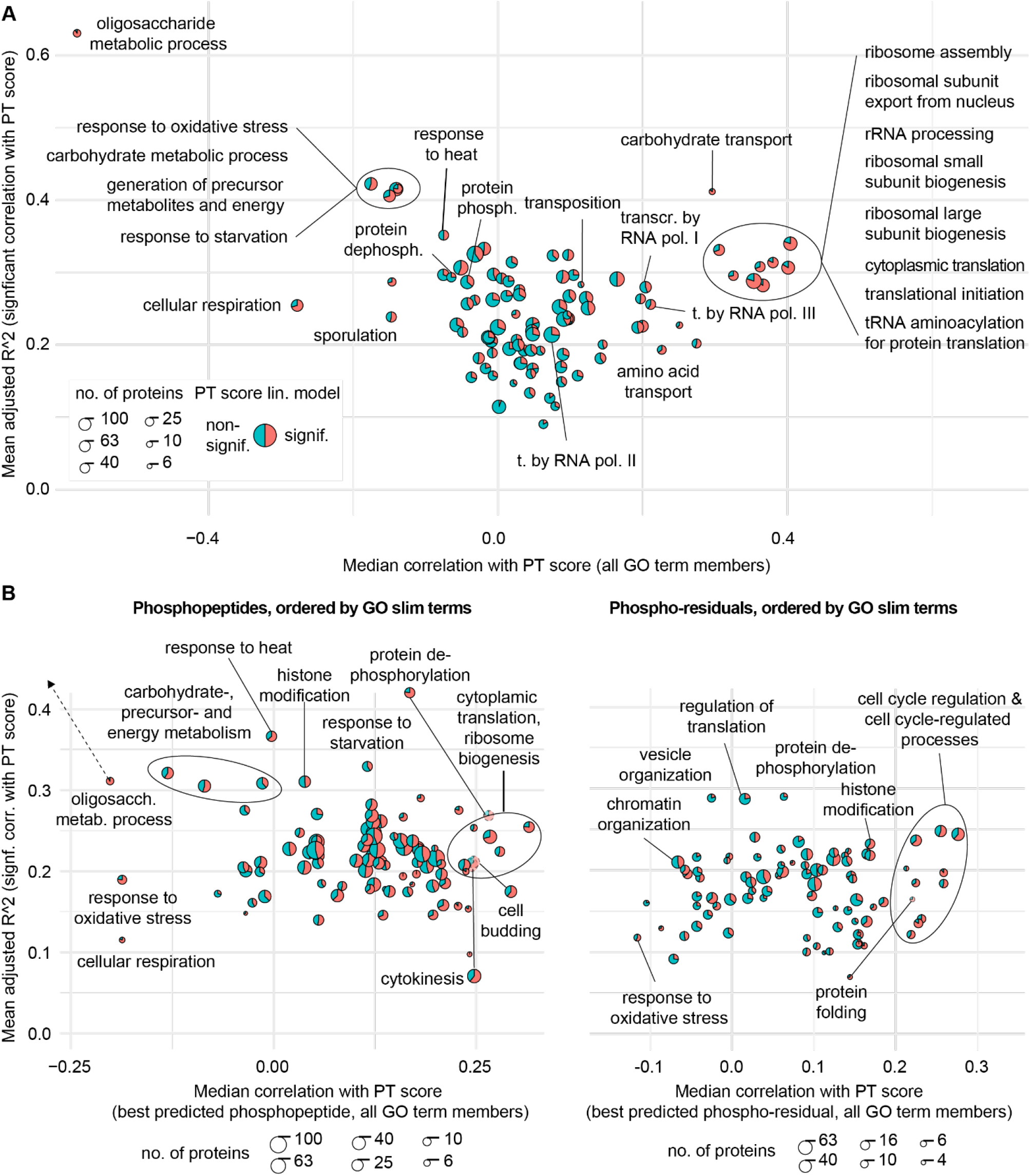
Correlation between PT score and functional modules (GO slim terms). **A** Illustration of linear model properties using the PT score as predictor for protein abundances, separated by slimGO terms. The size of each circle represents the number of proteins per GO term (logarithmic scaling) as indicated. Red and blue segments correspond to the proportion of proteins for which the PT score is or is not a significant linear predictor (q < 0.05), respectively. Diagram created using R package “scatterpie”. **B** Same representation of slim GO terms as in A but based on best predicted phosphopeptide (left panel) or phosphopeptide-residual (right panel) for each GO-annotated protein (see text for details). The GO term “oligosaccharide metabolic process” was moved inside the diagram boundaries for visibility. Some terms were moved outside of grouped term boundaries for clarity, as indicated.

We also analyzed the correlation of the PT score with our protein phosphorylation data sorted by GO slim terms (Fig. 3B). We have used two different measures to quantify protein phosphorylation: first the ‘raw’ abundance measurement of a phosphopeptide and second protein abundance-corrected residuals (‘phospho-residuals’, (Grossbach *et al*., 2022)). Whereas the former reflects a combination of protein abundance changes and changes in protein phosphorylation, the latter measure is corrected for changes in protein abundance (i.e. the residues of a linear model regressing phosphopeptide levels on corresponding protein levels per strain) and solely quantifies changes in phosphorylation. When several phosphopeptides were measured per protein, we considered the peptide that was best predicted by the PT score (highest adj. R^2^) to reflect the functional state of the corresponding protein (see Methods for a discussion of potential bias). The most positive correlations between the PT score and absolute peptide phosphorylation levels were again found in terms related to cytoplasmic translation and ribosome biogenesis, indicating concurrent abundance and phosphorylation increase in this process. Phosphopeptides from proteins involved in cell budding (mean adj. R^2^ = 0.21) and cytokinesis (mean adj. R^2^ = 0.21) were more strongly correlated with the PT score than the respective protein abundances (mean adj. R^2^ = 0.14 and 0.13, respectively). Consistently, we found that phospho-residuals belonging to proteins annotated with regulation of the cell cycle as well as processes known to show fluctuating phosphorylation through the cell cycle (Campbell et al., 2020), were positively correlated with the PT score. Interestingly, phospho-residuals of a number of proteins involved in dephosphorylation showed the highest mean correlation with the PT score. Finally, we noticed strong correlations between the PT score and phosphorylation events on enzymes involved in histone modification centered on H3K36 trimethylation, an important prerequisite for the expression of stress response genes (McDaniel and Strahl, 2017; Separovich et al., 2022). Together, these results revealed widespread adaptation of cellular processes coincident with changes in the PT network state reaching far beyond the core energy metabolism. Further details can be found in Supplemental Text 1.

### 6. A model for classification of genetic variant effects by network range

Our analysis of genetic variance in the BYxRM cross indicated that pQTL hotspots in particular were accompanied by significant PT score differences, resulting in a global adaptation of the molecular configuration of the cell. This led us to distinguish global (large-scale) network effects, like those induced by the *IRA2* locus, from local and regional effects.

Among loci with small differences in the PT score, some had detectable effects only on single proteins (Hsp104, Mog1), thereby remaining “local”, whereas others had effects on several or even many proteins without globally re-organizing the balance between diverse functional modules (which we call “regional” network effects). For example, strains with opposing alleles at the hotspot on Chromosome X (“ChrX:1”) did not show a significant PT score difference (−0.18, p = 0.5, Fig. S2D). Among the 20 proteins with a significant pQTL linking to this hotspot, several were reported to be targets of the zinc-responsive transcription factor Zap1 (Adh4, Tsa1 and Lap3 (Lyons et al., 2000; Wu et al., 2008)). Alleles of *ZAP1*, which coincides with the hotspot, contain 9 missense variants between BY and RM. The hotspot hence likely affected a functional module related to the sensing of zinc while the lack of a PT score difference indicated that PKA or TOR signaling were not affected. Further, in contrast to the *IRA2* locus we did not detect any large-scale reorganization of cellular process associated with the *ZAP1* locus. Thus, we classify the effect of the *ZAP1* locus as ‘regional’.

Our proposed classification of genetic variant effects into global, regional and local implies differences in terms of the distances between affected macromolecules across the network. To quantify this spreading of effects across the molecular network, we exemplarily computed pair-wise shortest path distances between all target proteins of the *IRA2* and *ZAP1* loci in a network of physical protein interactions (STRING, (Szklarczyk et al., 2021)) (Fig. 4A). Network distances between targets of the *ZAP1* locus were on average shorter (3.02 edges per pair, 153 pairs) than those between targets of the *IRA2* hotspot (3.46, 19503 pairs, p < 0.01 assuming a Poisson distribution for inter-node distances, Fig. 4A). Further, *ZAP1* targets comprised no “long-distance pairs” (here defined as pairs spanning more than 6 edges) whereas 3.7% of the *IRA2* target pairs were long-distance pairs. We conclude that effects of the *ZAP1* locus remained more confined or “regional” than those of the *IRA2* hotspot. Since the spread of the hotspot effect across the network might be related to effect strength or the number of targets, we investigated pair-wise distances for the targets of all 12 pQTL hotspots in the same manner as above and compared the mean distance of the target pairs and the fraction of long-distance pairs to the number of targets of the respective hotspot (Fig. S4A). This analysis did not support a correlation between effect spread and the number of targets. It further showed that distances between targets of the *IRA2* hotspot were greater than for other hotspots with comparable numbers of targets.

**Figure 4:**
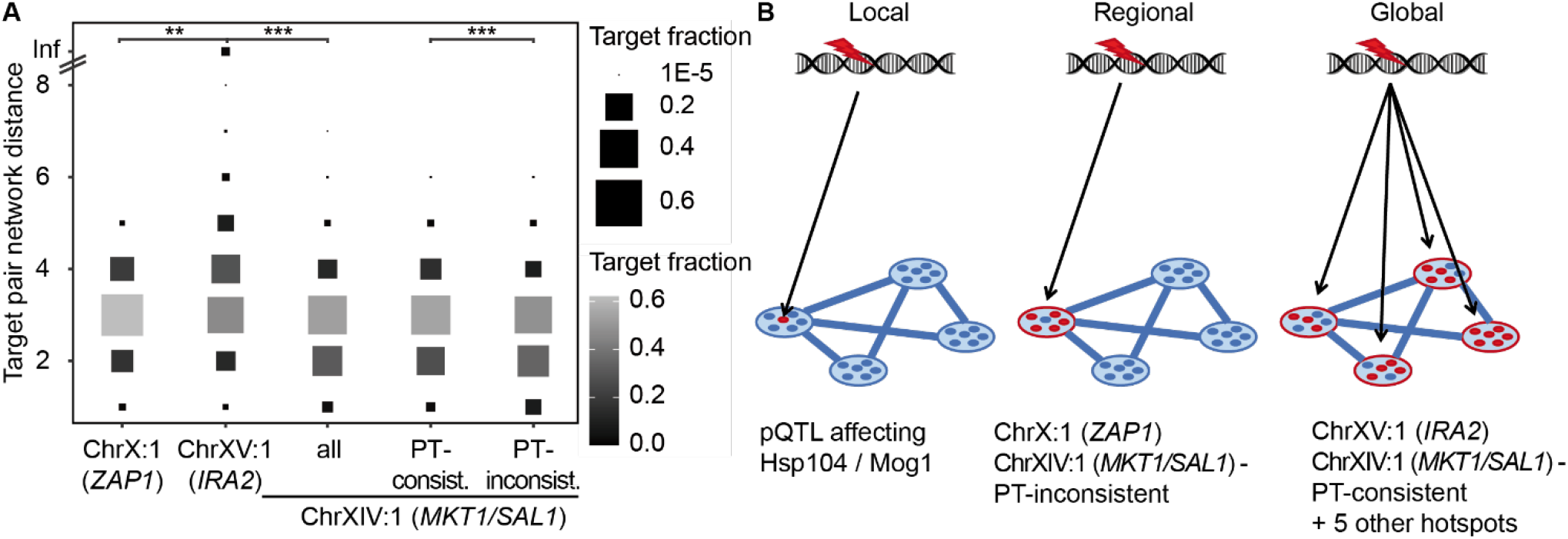
Model for classification of genetic variants according to their network effects. **A** Distribution of shortest path distances (STRING database physical interaction-based graph at medium confidence level (score > 0.400) for all pair-wise combinations of targets (or target subgroups as indicated) of the indicated pQTL hotspots. Some targets of pQTL hotspot ChrXV:1 (close to *IRA2*) could not be connected on this graph (indicated by a line at the top of the plot) and were assigned the highest distance of other pairs (8). **B** Proposed classification of genetic variant effects according to the network position of affected molecules.

We concluded that many but not all hotspots have a significant effect on the PT score. Substantial changes of the PT score (as exemplified by the *IRA2* locus) are associated with large-scale changes of the molecular network and a re-organization of many basic cellular functions, resulting in global network effects. Other hotspots, such as the *ZAP1* locus affect confined network regions and individual cellular functions, thus remaining regional.

The network effect of hotspot “ChrXIV:1” (coinciding with the *MKT1/SAL1* locus and with PTQTL2) was surprising. We expected to observe similarly far-reaching effects as for the *IRA2* hotspot due to the significant change in the PT score at this locus. However, pairwise network distances between protein abundance targets of hotspot ChrXIV:1 were on average shorter than for targets of any but one of the other hotspots (ChrXII:1, Fig. S4A). Protein abundance changes due to the RM allele at hotspot ChrXIV: 1 were globally anti-correlated with PKA and TOR inhibition (r = −0.44, p < 1e-27, Fig. S4B). Interestingly, we observed that mitochondrial proteins were not reduced as expected based on the PT score increase associated with the RM allele (Fig. S4C). Given the lower PT score in strains with the BY allele at this locus, we speculated that mitochondrial defects due to BY alleles of *MKT1* and *SAL1* (Dimitrov *et al*., 2009) led to reduced PKA or TOR activity following AMPK/Snf1 activation (Kingsbury et al., 2015; Malecki et al., 2020) or similar feedback mechanisms. Thus, we interpreted the effect of this hotspot as a combination of a regional effect on mitochondrial function and an additional effect via PKA or TOR signaling on a different set of cellular processes. To elucidate these effects, we partitioned the pQTL targets of this hotspot into two groups, depending on whether their abundance change was consistent (“PT-consistent”, 216 proteins) or inconsistent (“PT-inconsistent”, 210 proteins) with the difference in PT score due to the allele at this locus (Fig. S4D). Many (108/210) of the proteins in the PT-inconsistent group belonged to a limited set of functional modules including mitochondrial translation and electron transport chain function (Fig. S4C). Next, we calculated pair-wise shortest path distances across the STRING physical interaction network within the PT-consistent and -inconsistent groups. The average distance between PT-consistent targets was larger (2.85 edges) than the average distance in the PT-inconsistent group (2.61 edges, p < 0.001, Fig. 4A), suggesting that the former group of proteins belonged to more distant parts of the molecular network. Hence, we propose that the *MKT1/SAL1* locus had two distinct network effects: the first one resulting from long-range effects via the PT network state and a second one directly impinging on mitochondrial function without acting via PKA and/or TOR signaling.

Taken together, we were able to differentiate the effects of pQTL according to the relatedness or distance of their targets in a physical interaction network. Specifically, comparison of the proteome-wide effects of pQTL hotspots to changes in the PT network state allowed us to develop mechanistic hypotheses for the occurrence of global as compared to regional and local network effects (Fig. 4B).

### 7. Gene-environment interactions shape the PT network state

Previous studies have noted wide-spread gene-environment interaction for variants that influence gene expression in the BYxRM cross (Smith and Kruglyak, 2008). Gene-environment interactions were stronger for distant than for local QTL linkages, indicating that genetic effects acting in *trans* depend more strongly on the environment. The PT score is calculated from a number of different proteins, hence reducing the impact of individual *cis* effects and leading us to assume that changes are even more likely to occur via *trans* effects as compared to expression of an individual gene. To evaluate environmental influences on network effects of genetic variation, we investigated PT score changes in the BYxRM cross across two extreme growth conditions: growth on glucose against growth on ethanol.

First, we re-analyzed transcriptome data generated previously for strains of the BYxRM collection grown on glucose and on ethanol (Smith and Kruglyak, 2008). In this dataset, 39 induced and 28 repressed PT marker transcripts among the sets derived from chemical inhibition data (see Methods) were present and could thus be used to calculate a PT score for each of 109 strains in both conditions. Across 99 strains for which proteome data was available in our own dataset, PT scores for strains grown on glucose showed good agreement between the two datasets (Spearman’s ρ = 0.57, p < 1E-9, Fig. 5A), confirming that the PT score was consistent across independent studies and different molecular modalities. Notably, PT scores of strains grown in glucose and ethanol were mostly positive and negative, respectively, indicating a strong effect of the growth condition (Fig. 5B), while PT scores of the same strains were correlated between the two conditions (ρ = 0.52, p < 1E-9). Further, the transcriptome variability was correlated with the PT score under both growth conditions: PCA analysis of transcriptomes across 109 strains revealed a strong correlation between PC1 scores and PT scores in glucose (adj. R^2^ = 0.91, Fig. 5C). The correlation was slightly reduced for strains grown on ethanol (adj. R^2^ = 0.86) and PC1 reflected a smaller proportion of total transcriptome variance (Fig. 5C), in line with the PT scores spanning a smaller range compared to the glucose condition (Fig. 5B). While this was consistent with lower activity of the PKA and TOR pathways on non-fermentable carbon sources, our results indicate that the PT network state dominated the molecular variation between strains even during respiratory growth.

**Figure 5:**
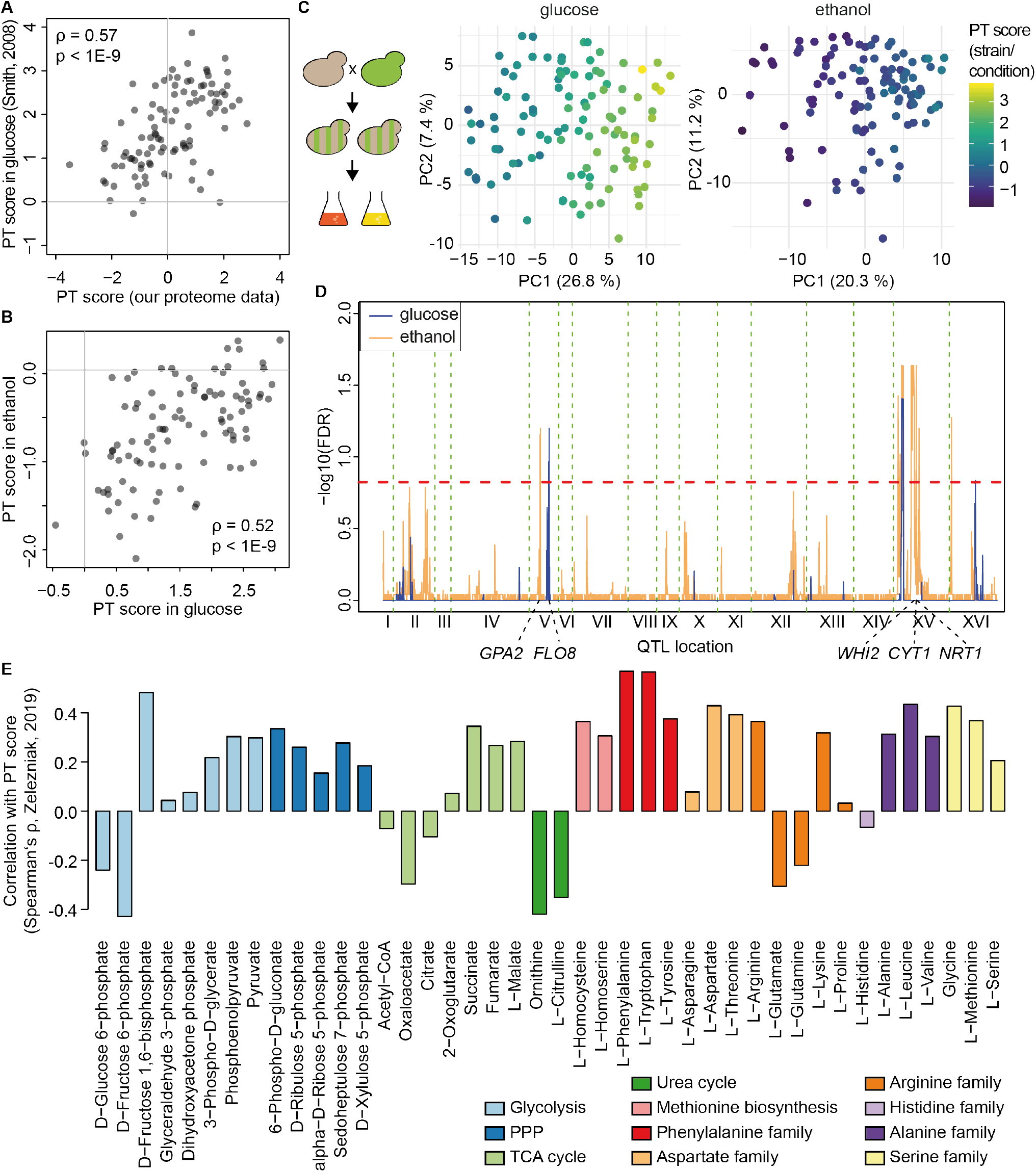
The PT network state is subject to gene-environment interaction. **A** Comparison of PT score based on our proteome data and PT score based on transcriptome data from (Smith and Kruglyak, 2008) for 99 strains grown on glucose and present in both datasets. **B** Distribution of PT scores for 109 strains grown in glucose (left) or ethanol (right) based on transcriptome data from (Smith and Kruglyak, 2008). **C** PCA of transcriptome variability for 109 strains grown in glucose or ethanol as indicated. Calculation of Eigenvalues, coloring by PT score and labeling as in Fig. 2A. **D** QTL mapping of PT score for 99 strains grown in glucose (blue) and ethanol (orange). 15 % FDR threshold indicated by dashed red line. **E** Correlation of PT score based on proteomic data for several kinase KO strains with metabolite measurements (Zelezniak et al., 2018).

We next performed QTL mapping for PT scores of strains grown on glucose and ethanol as above. The locus close to the *IRA2* hotspot harbored a significant PTQTL in both conditions (Fig. 5D). There was a significant negative interaction between growth condition and the allele at this locus (−0.48, p < 0.01 for the interaction term in a combined linear model), indicating that the effect was significantly weaker during growth on ethanol. This is consistent with the observation that fewer transcripts linked to this region for strains grown on ethanol (Smith and Kruglyak, 2008). Another PTQTL for strains grown on glucose was found within 3 kb to the *FLO8* gene on Chromosome V, which encodes a transcription factor involved in filamentation and that carries a null mutation in the BY progenitor strain S288c (Liu et al., 1996). Several PTQTL for strains grown on ethanol were close to genes that are known to be relevant for PKA or TOR signaling, especially in conditions of nutrient scarcity (Fig. 5D). These include the *GPA2* gene on Chromosome V, which has recently been associated with a number of cellular fitness traits in this cross (Nguyen Ba *et al*., 2022). Comparison of a region harboring three PTQTL on Chromosome XV to the same fitness dataset indicated that variants with pleiotropic effects in *WHI2*, *CYT1* and *NRT1* might be causal for PT score differences in ethanol. Consistently, all of them are involved in processes required to cope with nutrient scarcity. Whi2 negatively regulates TORC1 and Ras/PKA activity under nutritional stress (Chen et al., 2018; Leadsham et al., 2009; Teng and Hardwick, 2019). The gene product of *CYT1*, cytochrome C1, is essential for respiration and finally, *NRT1* encodes a high-affinity nicotinamide riboside transporter (Belenky et al., 2008). In sum, this analysis demonstrated that the PT score correlated with molecular network effects of genetic variation in multiple growth media in a condition-dependent manner and that large-scale transcriptome variation was associated with the PT score under those conditions.

### 8. The PKA/TOR network state correlates with an anabolic pattern in kinase KO strains

Our analysis of proteomic changes associated with PT state differences indicated strong links to the central energy and amino acid (AA) metabolism (Fig. 3A), which aligns with the known functions of PKA and TOR signaling. PKA activity is subject to sensing of external and internal metabolic cues, most notably the levels of glucose and cAMP, respectively (Conrad *et al*., 2014), whereas the activity of TOR is strongly dependent on the sensing of intracellular and extracellular AA levels (Gonzalez and Hall, 2017; Hara et al., 1998; Shimobayashi and Hall, 2016). In order to further investigate associations between the PT network state and the metabolic configuration of the cell, we re-analyzed proteomic and metabolomic data for kinase knockout strains (KOs) (Zelezniak *et al*., 2018). We focused on metabolite data recorded for 22 kinase KO strains (datasets 2 and 3 in (Zelezniak *et al*., 2018)) and compared PT scores calculated based on proteomic data to metabolite levels (Fig. 5E). We observed that 15 out of the 18 proteinogenic AAs measured showed positive correlation with the PT score, with notable exceptions being the levels of glutamate (Glu, ρ = −0.31) and glutamine (Gln, ρ = −0.22). This may seem surprising, since addition of Gln promotes strong activation of TOR signaling (Duran et al., 2012; Oliveira et al., 2015). However, our analysis differs from studies employing the addition or removal of extracellular Gln as a nitrogen source in that it investigates the metabolic state across strains in which PKA and TOR activity differ due to genetic perturbation. Since Glu and Gln are essential nitrogen donors for the synthesis of both nucleotides and AAs, they will be rapidly used in highly proliferating cells. Thus, in this setting, intracellular concentrations of Gln and Glu might be lower in strains with high PT scores due to more active proliferation. Our observation is also consistent with earlier reports that TOR activity represses Glu and Gln biosynthesis via inhibition of the retrograde signaling pathway (Dilova et al., 2004) perhaps because high levels of other AA reduce the need for transamination from Glu and Gln. Negative correlation of the PT score with the AA precursor oxaloacetate (ρ = −0.30), no correlation with 2-oxoglutarate (ρ = 0.07), and positive correlation with pyruvate (ρ = 0.30) indicated that glycolytic activity was the primary source of alpha keto acids in strains with high PT activity. A particularly strong correlation was observed between the PT score and fructose-1,6-bisphosphate (ρ = + 0.48), which is in line with the decisive regulatory role of this metabolite for Ras/PKA and as an indicator of glycolytic flux (Peeters et al., 2017; Tanner et al., 2018; Zhang and Cao, 2017). Finally, we observed positive correlation of the PT score with all intermediates of the pentose phosphate pathway (ρ = + 0.15 to ρ = + 0.28). In sum, our analysis confirmed a tight association between the PT network state and cellular metabolism. In particular, PT activity seemed to be linked to an anabolic and proliferative state of the cell.

### 9. The PKA/TOR network state explains molecular and phenotypic diversity in distantly related yeast species

We next asked whether a similar network state exists in different species and focused first on the distantly related fission yeast *Schizzosaccharomyces pombe*. We previously observed differences in stress resistance and longevity between a natural isolate, Y0036 (Y0) and the common *S. pombe* lab strain Leupolds968 (L9) (Clement-Ziza et al., 2014). We also noticed increased stress resistance across several conditions in the industrial strain DBVPG2812 (DB, unpublished observation). To explore the genetic underpinning of that phenotypic diversity, we established a three-way cross between these parents (R1 = Y0 x L9, R2 = L9 x DB and R3 = DB x Y0) with 43-45 segregants per cross and collected transcriptomic data for the parental strains and each segregant during exponential growth on standard medium. Furthermore, we also collected these data for populations grown in the presence of 0.5 mM hydrogen peroxide (H_2_O_2_) for 1 hour. Oxidative stress constitutes a naturally occurring stress that plays a decisive role for the network state re-organization at the transition to respiratory growth (Tran et al., 2019).

The PT score for each segregant strain was calculated based on orthologs of the budding yeast PT score marker genes, during both unperturbed growth and in the presence of H_2_O_2_. The range of the PT score varied strongly between the individual pair-wise crosses (R2 > R1 ≈ R3). Segregants in the R2 cross were assigned the highest median PT score (1.57) in the unstressed condition but the lowest median score (−1.27) in the H_2_O_2_ stress condition (Fig. 6A). As previously observed in budding yeast, the PT score correlated with segregant scores along PC1 of the transcriptome in both conditions (adj. R^2^ = 0.72 and 0.82, both p <1E-15, Fig. 6B).

**Figure 6:**
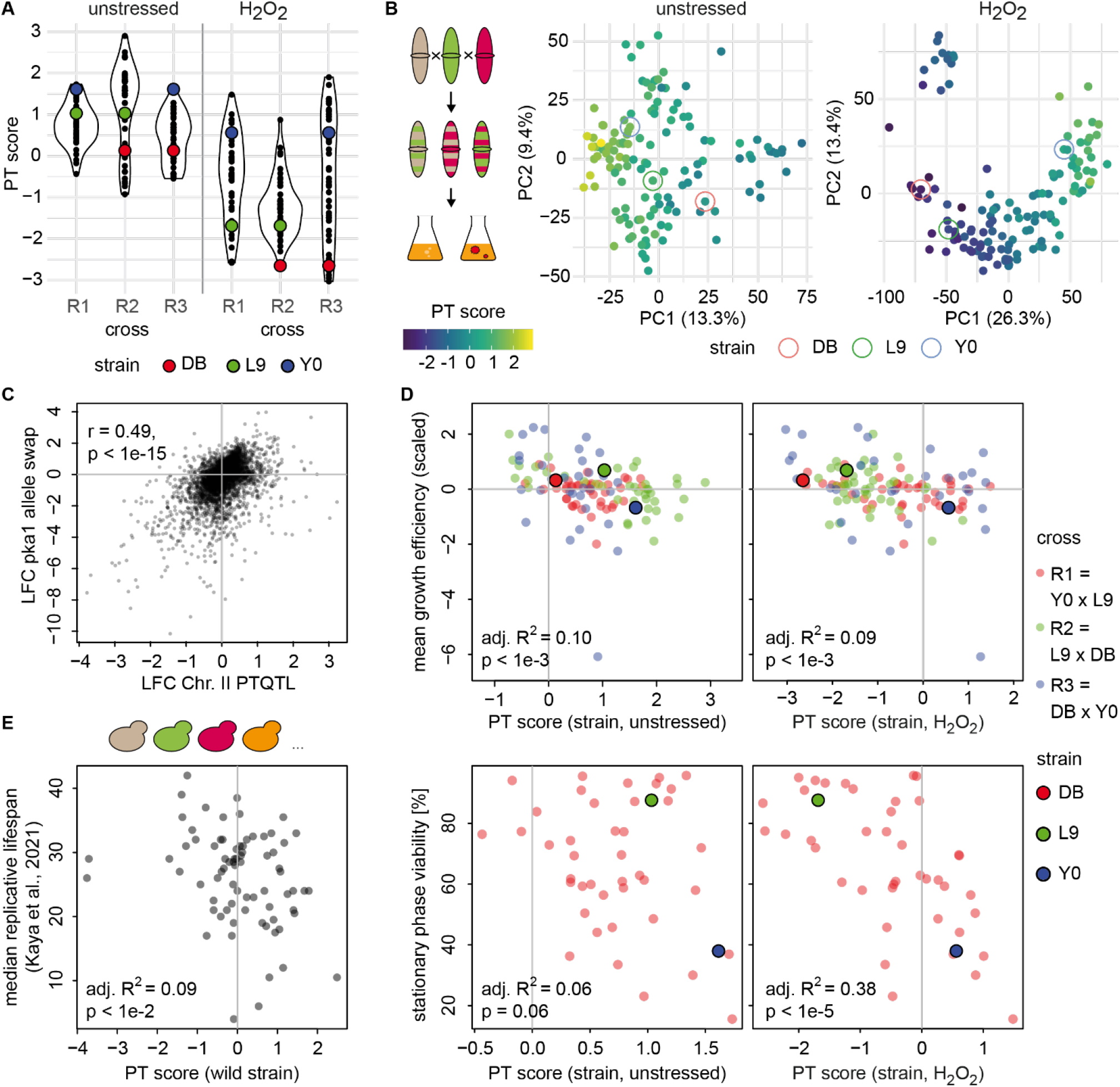
PT score explains molecular variability and cellular fitness traits in distant yeast species. **A** Range of PT score values across pair-wise crosses (R1 = Y0 x L9; R2 = L9 x DB; R3 = DB x Y0) and in two conditions. Parental strains highlighted as indicated. **B** PCA based on transcriptome variability for 127 segregants and 3 parental strains as indicated, in unstressed (left) or H_2_O_2_-stressed condition (right). Coloring by PT score and labeling as in Fig. 2A. **C** Comparison of pka1 DB-allele effect in an L9 allele-swapped strain to differences due to the DB allele at the PTQTL on Chromosome II in the 3-way cross for all transcripts detected in both settings. **D** Correlation between growth efficiency and PT scores for 114 strains (upper panels) and comparison between stationary phase viability and PT scores for strains in the R1 cross (lower panels). PT scores were determined for samples of each strain grown either in the presence or absence of H_2_O_2_. Coloring by cross and parental strains enlarged as indicated. **E** Comparison between median replicative lifespan and PT score for 75 wild isolate yeast strains based on data from (Kaya *et al*., 2021).

Next, we performed QTL mapping to identify genetic determinants of the PT network state in fission yeast. In the unstressed condition, the strongest QTL was detected at an FDR of 0.15 (Fig. S5A). We investigated this locus and found a polymorphism at position 1073 in the *pka1* allele of the DB parent that led to an amino acid exchange from cysteine to phenylalanine (C358F) compared to the other parents. Indeed, swapping the *pka1* allele in the lab strain L9 for the DB allele resembled the effects of this locus on the transcriptome (r = 0.49, p < 1E-15, Fig. 6C). We still observed strong correlation between the PT score and segregant scores along PC1 in the group of segregants with the DB allele (adj. R^2^ = 0.63, p < 1E-8). Hence, additional genetic variants likely contributed to variation of the PT network state in the unstressed condition but were beyond our limit of detection. QTL mapping in the H_2_O_2_ stress condition revealed significant signals in six distinct regions (Fig. S5A). One of these QTL coincided with the *pka1* locus and consistently, the *pka1* allele swap also partially reproduced the effects of this QTL across the transcriptome in the presence of H_2_O_2_ (R = 0.25, p < 1E-15). The strongest signal for PT score changes in H_2_O_2_ was attributed to a region on Chromosome I, within a large genomic inversion in the Y0 parental strain.

The Y0 allele at the most significant marker led to higher PT scores in the presence of H_2_O_2_ (+1.43, p < 1E-6), but not during unperturbed growth (+0.13, p = 0.4) and contributed to the dominance of PT score-related differences in global transcriptome variability of R1 segregants specifically in the H_2_O_2_ condition (Fig. S5B). This effect again illustrated the importance of gene-environment interactions underlying PT network state differences.

To evaluate the relevance of the PT score for cellular fitness in fission yeast, we recorded growth efficiency (final OD after 36 h of growth in unperturbed conditions) for 114 strains across the collection. This measure correlated significantly with both the strains’ PT scores in unperturbed growth conditions as well as with PT scores for strains grown in the presence of H_2_O_2_ (Fig. 6D, adj. R^2^ = 0.10 and 0.09, respectively, both p < 1E-3). We further evaluated the predictive capacity of PT scores in fission yeast using stationary phase viability measurements obtained in a previous study for the R1 cross (Clement-Ziza *et al*., 2014). As for the previous trait, we compared these measurements to PT scores from the unperturbed and oxidative stress conditions. In contrast to growth efficiency, we observed a strong negative correlation between stationary phase viability and the PT scores calculated from populations grown in the presence of H_2_O_2_ (adj. R^2^ = 0.38, p < 1E-5) but only weak correlation with the PT scores based on unperturbed samples (adj. R^2^ = 0.06, p = 0.06). As described above, we detected a major QTL that dominated PT score differences among strains in the R1 cross exclusively in the presence of H_2_O_2_. We speculate that such condition-specific effects of genetic variance can contribute to fitness differences by eliciting PT network state differences. Such effects would be limited to fitness traits that are related to the conditions in which the genetic variant effects are penetrant.

Motivated by the successful application of the PT score as a predictor of cellular fitness traits in fission yeast, we applied it to replicative lifespan measurements recorded for 75 wild isolate strains of the yeast species *Saccharomyces paradoxus* and *S. cerevisiae* (Kaya et al., 2021). Based on the same sets of markers derived from chemical inhibition experiments as before, we calculated PT scores using the reported transcriptomic data and compared these scores to the replicative lifespan of the corresponding strains (Fig. 6E). Indeed, PT scores were weakly, but significantly predictive of replicative lifespan (adj. R^2^ = 0.09, p < 0.01) across these wild isolates.

We conclude that the PT network state explains long-range network effects and fitness consequences of genetic variants across diverse growth conditions and large evolutionary distances.

## Discussion

In this study we have investigated adjustments of ‘multi-omics’ molecular network states as a result of genetic variation in three yeast species during exponential growth across different environmental conditions. Our work represents the first systematic investigation into the propagation of natural genetic effects across large distances within a molecular network and provides a mechanistic explanation for a range of perturbations that gave rise to such ‘global’ effects. We observed that substantial fractions of transcriptome, proteome and phospho-proteome variability among genetically different yeast strains were correlated to a score based on markers of PKA and TOR pathway activity (PT score). This strong dependency was evidenced for example by the fact that the first components of respective PCAs strongly correlated with the PT score, indicating that the underlying regulatory program ‘moves’ large parts of the molecular network along a single-dimensional axis.

PKA and TORC1 signaling pathways regulate the activity of a wide range of cellular functions beyond growth, intermediate and energy metabolism, including processes such as transcription, mitochondrial and cytoplasmic translation, spore formation and autophagy in yeast and other organisms (Conrad *et al*., 2014; Gonzalez et al., 2020; Plank, 2022; Workman et al., 2016). The roles of PKA and TORC1 seem to be rather temporally than functionally distinct (Kunkel *et al*., 2019). We did not specifically investigate the role of TORC2 signaling for the program reflected by the PT score but do not exclude a potential influence, since TORC2 was recently shown to share common targets with PKA and TORC1 via its downstream kinase Ypk1 (Plank et al., 2020). Other pathways, including AMPK/Snf1 signaling, regulate overlapping processes and cross-talk with PKA and TOR but affect limited sets of proteins if modulated independently (Kingsbury *et al*., 2015; Malecki *et al*., 2020; Zaman et al., 2009).

The PKA and TOR pathways jointly coordinate the transition from fermentative to respiratory metabolism at the diauxic shift (Pedruzzi *et al*., 2003). Earlier studies have already provided examples for genetic variants that affect this metabolic balance, including mutations in fission yeast pyruvate kinase (Kamrad *et al*., 2020), but have not systematically explored its genetic basis. Our QTL mapping of the PT score revealed that tuning the state of gene expression in a manner similar to the diauxic shift is a polygenic feature in exponentially growing yeast. Clearly, yeast cells can adopt metabolic states on a continuous spectrum between opposite poles – rather than switching between purely fermentative and purely respiratory metabolic states. By application of the PT score in different growth conditions and across an ensemble of wild yeast strains, we were further able to show that gradual variation of the underlying gene expression program was not an artefact of a particular yeast cross or individual mutation. Therefore, we propose to think of PKA and TOR-related gene expression as an example of a widely observable ‘cellular network state program’ (here, the PT network state).

We were able to discriminate between loci that caused large-scale (virtually global) changes of a broad network state and loci that had more confined consequences on smaller parts of the molecular network. Examples for the latter type of loci included pQTL with local effects on the abundance of proteins associated with thermotolerance (Mog1, Hsp104) and a pQTL hotspot close to *ZAP1* with regional effects on several downstream targets of the Zap1 transcription factor but without inducing a large-scale reorganization of multiple functional modules. Most hotspots of protein abundance regulation coincided with detectable changes in the PT score. Such ’global effect variants’ led to altered expression of many proteins in a manner that was largely consistent with changes to PKA and/or TOR activity. Our analysis of the association of different cellular functions (GO terms) with the PT network state at the proteome and phosphoproteome level confirmed the coordinated adaptation of a wide range of functional modules to the metabolic state of the cell. Importantly, here we did not study the targets of a specific pathway or perturbation but rather correlations with a global network state, shifted by natural genetic variation. Our work suggests that the PT network state mediates the effects of genetic variation on a wide spectrum of cellular traits, including growth on different carbon sources, stress response (heat stress, oxidative stress) and replicative lifespan. This provides a new perspective on the integration of metabolic regulation with other cellular functions in the context of genetic variability, which has become a major focus of studies in the field of proteostasis and ageing (Ottens et al., 2021).

Our analysis framework and the concept of molecular network states that we are proposing helps to better understand (i) the polygenicity of complex traits, (ii) the propagation of genetic effects in molecular networks and thereby (iii) pleiotropic effects of single variants. Regarding the first point, we have shown that the PT network state better predicts the heat-induced lag trait than any individual QTL and adds predictive power to a model containing all significant QTL in our study. Thus, the PT score is an integrative measure of a molecular network state that combines the influences of numerous genetic variants. Regarding the second point, our distinction of local, regional and global network effects provides a conceptual model for characterizing systems-level consequences of genetic variants. We propose that far-reaching genetic effects, such as those postulated in the concept of ‘omnigenic inheritance’ (Boyle *et al*., 2017) may originate from global adjustments in cellular physiology, such as transitions from fermentative to respiratory energy metabolism. Consequently, pleiotropic effects of genetic variants are conditional on the network regions and functional modules affected by them. Variants altering the global balance between functional modules will affect a broad spectrum of cellular traits.

PKA and TOR signaling have been established as central switches in cellular physiology across most eukaryotes (Gonzalez *et al*., 2020). Exemplarily, we confirmed conservation of the PT network state between *Saccharomyces cerevisiae* and *Schizzosaccharomyces pombe* despite millions of years and different paths of evolution that separate these species (Rhind et al., 2011). We propose that similar global network state changes may be observable in mammalian cells, for example during the transition between respiratory and glycolytic metabolism. This transition is known as the ‘Warburg effect’ in cancer (Liberti and Locasale, 2016) but has been described in many other circumstances and was recently proposed as a pathogenic mechanism in sporadic forms of Alzheimer’s disease (Traxler et al., 2022). Further, transitions between these two metabolic states seem to be associated with cellular differentiation processes (Bhattacharya et al., 2020; Zhang et al., 2018). Our work does not exclude the possibility that signaling pathways other than PKA (ERK) and mTOR play a similar globally coordinating role in metazoan cells. We have shown the utility of quantifying network states in the study of cellular traits including longevity, oxidative stress and thermotolerance, which are important models for complex human diseases. Thereby, network states such as the one presented here may help to explain a relevant proportion of the polygenicity of complex traits in higher eukaryotes.

## Acknowledgements

MW and JG acknowledge funding by the Deutsche Forschungsgemeinschaft (German Research Foundation, DFG), Grant-IDs: 440376057 (MW), CRC 1310 325931972 (JG).

## Methods

### Heat-stress experiments

The BYxRM yeast strain collection was originally derived from a cross between the two parental strains BY4716, an S288C derivative (*MAT*α lys2Δ0) and RM11-1a (*MAT*a leu2Δ0 ura3Δ0 ho::KAN)(Brem *et al*., 2002). Experiments to determine thermotolerance were performed for a subset of 100 segregants, which were selected based on non-flocculent growth and by the availability of genotype data from our previous study (Grossbach *et al*., 2022), and for the parental strains. Experiments on the parental strains were performed alongside with segregant experiments on three separate days to assure consistent results over the entire course of the experiments. Pre-cultures in synthetic complete medium with 2 % glucose (FORMEDIUM) were inoculated from YPD plates grown for 3 days after thawing of yeast stocks. Over-night cultures in the same medium were inoculated from the pre-cultures and were grown with constant shaking at 30 °C in glass flasks. Heat-treatment was applied to samples of cells taken at OD_600_ between 0.7 and 0.8 in PCR tubes in a PCR cycler. Temperature was increased from 25 °C to 45 °C by 0.1 K/s with subsequent incubation at 45 °C for 8 min. As a control, samples from the same cultures were incubated at 25 °C. 60 μl of the samples were then diluted 1:30 into fresh medium in 24-well plates. Growth curves following dilution after mock or heat treatment were recorded in a Thermo Fisher Varioskan Flash instrument with further incubation and shaking at 30 °C. Growth curves were fitted using the R package ‘grofit’ (Kahm et al., 2010) with a parametric fit following a Richard’s law model to infer growth characteristics.

### QTL mapping and trait mapping based on molecular data in budding yeast

QTL mapping to identify genetic determinants of heat-induced lag and the PT score in budding yeast was performed using Random Forest (RF)-based machine learning as described previously (R package “RFQTL” and (Grossbach *et al*., 2022)). We used a similar procedure to determine protein abundances that have predictive power for heat-induced lag. In brief, scaled protein abundance values were used as continuous predictor variables to grow RF models of heat-induced lag measurements from 102 strains, including averaged triplicate measurements for the parental strains. Statistical significance of the results was assessed by permutation of heat-induced lag measurements across the strains to generate a null distribution of variable importance for each predictor. P values were adjusted for multiple testing using a Benjamini-Hochberg procedure.

QTL mapping of the PT score based on data reported in (Smith and Kruglyak, 2008) was performed on a set of 99 segregants, which we previously genotyped based on our own RNA-Seq data (Grossbach *et al*., 2022).

### PT score calculation

We defined sets of PT-regulated transcripts by setting a threshold of at least two standard deviations for the effects of chemical inhibition (Kunkel *et al*., 2019). Here, we included data at both the 20 min and 150 min time-points for both PKA and TOR inhibition. After exclusion of transcripts that changed abundance in the opposite direction in any of these treatments, we retained 47 (of 65) transcripts as a PT-induced set and 44 (of 90) transcripts as a PT-reduced set. In our data, abundance measurements for 22 and 18 corresponding proteins was available. A PT score was then calculated for each of 112 strains for which proteomic data was available (Grossbach *et al*., 2022) as the difference between the median abundance of proteins in the induced set to the median in the reduced set using protein abundances after normalization and centering across all strains. Thus, a higher PT score corresponds to higher median abundance of PT-induced proteins relative to PT-reduced proteins. The same procedure was applied to calculate PT scores for 109 segregants based on transcript abundance data reported in (Smith and Kruglyak, 2008), for 96 kinase KO strains based on proteome data reported in (Zelezniak *et al*., 2018) and for transcriptome data for 133 strains from our three-way cross in fission yeast.

### Analysis of phospho-proteome correlations

Since in many cases, several phosphopeptides were measured per protein, we considered the peptide that was best predicted by the PT score (highest adj. R^2^) to reflect the functional state of the corresponding protein. This decision introduces a bias for proteins with high numbers of phosphopeptides and might in some cases overestimate the “net correlation” between a specific protein and the PT score. To mitigate the second and potentially more severe bias we tested each protein for inconsistent predictions by the PT score. For 97 proteins that contained more than one significantly predicted peptide (q < 0.05), inconsistent predictions were found in only 13 cases (13.4 %). This fraction was only slightly affected by the choice of the FDR threshold (15.8 % inconsistent among 120 proteins at q < 0.25). We did not exclude these proteins from the following analysis, since they are unlikely to affect correlations across entire GO terms.

### Generation of a three-way fission yeast cross

Parental strains for the three-way cross in fission yeast were chosen based on differences in their resistance to oxidative stress in preliminary experiments. The three parental strains were JB50, a strain closely related to the reference strain JB22 (Leupolds968, L968), JB759 (Y0036), a strain with increased sensitivity to oxidative stress compared to JB50, and JB760 (DBVPG2812), a strain that is more resistant to oxidative stress than JB50. We used a double selection strategy to obtain diploid hybrids between each pair of parental strains. Specifically, we deleted the *ade6* locus in each parental strain and differentially replaced it with resistance genes for either kanamycin (KAN, in JB50) nurseothricin (NAT, in JB759) or hygromycin B (HB, in JB760). Plates containing both fungicides corresponding to a pair of parental strains were used to select for diploid hybrids carrying both resistance genes. F1 haploid recombinants were obtained straight from the selected diploid hybrids by tetrad analysis. F2 haploid recombinants were obtained by performing tetrad analysis of F2 diploid hybrids obtained from a mass mating among F1 haploid segregants. The cross included 150 strains in total.

### RNA-seq analysis of fission yeast strains

Samples for transcriptomic analysis were grown in 50 ml YES medium to an OD595 of 0.4-0.5 at 32 °C. These samples were either harvested before or after the exposure to 0.5 mM H_2_O_2_ for one hour. In total, the transcriptomes of 286 samples, corresponding to 130 strains in two conditions were quantified. RNA isolation, library preparation and sequencing were performed as described previously (Clement-Ziza *et al*., 2014). We mapped reads against the Schizosaccharomyces pombe reference genome using Bowtie with the following parameters: -C -n 3 -e 100 -best (v.0.12.7, Langmead et al. (2009)). Read group information was added, and BAM files were sorted using Picard utilities (http://broadinstitute.github.io/picard). Further processing of the RNA-seq data was performed with the GATK pipeline (v3.4-46-gbc02625), according to best practice guidelines (Van der Auwera et al., 2013). Variants at the polymorphic sites were then called with UnifiedGenotyper. If the GATK score for a given site was below 20, the position was considered as unknown. We excluded polymorphic sites for which (i) the samples of the parental strains could not be called, (ii) or more than 50% of the segregants could not be called, or (iii) the minor allele frequency was less than 10% in our cross. We also excluded genetic markers if they were called differently than both closest neighbors (i.e. the nearest upstream marker and the nearest downstream marker) and those markers were within 50 kb. When possible, missing genotypes were inferred through the neighboring markers if they had a identical segregation patterns and were within 50 kb. In these cases the marker was assigned the same allele as the two flanking markers. VCFs were converted to strain-specific FASTA-files using the GenomeGenerator tool (Clement-Ziza et al., 2014).

For quantification, We mapped reads with Bowtie (v.0.12.7, Langmead et al. (2009)) against the strain-specific genomes we generated based on the RNA-seq data using the following options: - C --best -m 1. Genes to which no reads were mapped in at least 50% of samples were dismissed. To correct for differences in library size, we compute a correction factor for each sample. We determined the 20% and 80% quantiles for each sample i and computed the median Mi for all counts between these quantiles. The counts for gene j and sample i were then multiplied with the ratio of the median of the counts for this subset of genes and the mean of these medians across all samples to correct for different library sizes. For replicate measurements, counts were averaged.

### Growth profiling of fission yeast strains

Growth profiling of 150 strains of the three-way fission yeast cross was performed using the bioLector system (m2p-labs, Germany) as described previously (Clement-Ziza *et al*., 2014). The growth of each sample was determined by light scattering in 3-minute intervals for at least 25 hours and then converted to optical densities with a linear model. Growth efficiency was calculated as the difference between the initial and final OD. The strains were distributed over 28 batches. We corrected for batch effects by subtracting the mean of all measurements for the batch from each of these measurements. Afterwards, we employed a step-wise procedure to remove batches that showed much more variation than the rest of the batches. First, we computed the variance of all trait values per batch. Second, if any batch had a variance that was 2.5 times as large as the average variance of all batches, we removed the batch with the highest variance. Then we repeated this step until the variance for no batch exceeded the set threshold above the variance of the remaining batches. We removed four batches with this procedure, resulting in at least one measurement of growth efficiency for 114 strains. Repeated measurements per strain were averaged for analysis.

### Generation of a pka1 allele-replacement strain in fission yeast strain JB50

To investigate the effects of the polymorphism in pka1 (C358F between JB50 and JB760) we generated an allele replacement strain. This strain was identical to JB50 aside from position 1073 in the coding sequence of pka1, changing the respective codon from UGU, coding for cystein, to UUU, coding for phenylalanine. The strain was generated using a CRISPR/Cas9 method as described before (Rodriguez-Lopez et al., 2016) using the primer pair 5’- acataacctgtaccgaagaaAGCAACTGTTGTACTCTTTGgttttagagctagaaatagc-3’ (forward) and 5’- gctatttctagctctaaaacCAAAGAGTACAACAGTTGCTttcttcggtacaggttatgt-3’ (reverse). The replacement strain is referred to as PKA1Rep in the following. We used Sanger sequencing to validate successful introduction of the JB760-allele of pka1 in the JB50-background. To measure the effects of the pka1 allele-replacement on the transcriptome and proteome, we grew three replicates each of JB50 and PKA1Rep under normal growth conditions and three replicates, for each strain, which were exposed to increased oxidative stress (0.5 mM H_2_O_2_) for one hour before the samples were harvested. Each sample was separated into a fraction for RNA extraction and one for protein quantification. The fraction for protein quantification was centrifuged and the pellets washed with cold PBS and snap-frozen in liquid nitrogen for transport.

### Transcriptome quantification for the JB50 pka1 replacement strain

RNA was extracted with the hot phenol method described in (Lyne et al., 2003). RNA was further purified with Qiagen RNAeasy columns, and DNAse treatment was performed in the columns (as suggested by manufacturer) prior to library preparation. RNA quality was assessed with a Bioanalyzer instrument (Agilent, United States), and all samples had a RIN (RNA Integrity Number) > 9. cDNA libraries were prepared with the Illumina TruSeq stranded mRNA kit, according to the manufacturer’s specifications, by the Cologne Centre for Genomics (CCG) facility. The samples were sequenced on a single lane of an Illumina Hiseq4000 to produce stranded 2×75 bp reads. Reads were trimmed with Trimmomatric (v0.36 (Bolger et al., 2014)), with the following parameters differing from default settings: LEADING:0 TRAILING:0 SLIDINGWINDOW:4:15 MINLEN:25. The reference genome was indexed with bowtie2-build with default settings. Paired reads were aligned to the reference genome using bowtie2 with default settings (v2.3.4.1 (Langmead and Salzberg, 2012)). In the case of the allele replacement strain, the reference genome was edited to reflect the base substitution within pka1. Aligned reads were counted using intersect from the bedtools package (v2.27.1 (Quinlan and Hall, 2010)), with the parameters -wb -f 0.55 -s -bed. Identical reads were only counted once. Readcounts were tested for differential expression between strains using DESeq2 v1.18.1 with default settings (Love et al., 2014). We tested differential expression between strains separately with or without addition of H_2_O_2_.

## Supplemental text 1

We conducted an analysis of correlations between the PT score and proteins sorted by GO slim terms to leverage the availability of extensive -omics data on the collection of yeast strains derived from the BY and RM parental strains. The analysis charts the wide-spread adaptation of the molecular network along the axis defined by the PT score, which we consider to represent a fundamental adjustment of the network between the poles of fermentative versus respiratory metabolic preference. Consistently, cellular respiration was among the terms that contained proteins with predominantly negative correlations with the PT score. However, the average predictive accuracy of the PT score for the abundance of these proteins was less than in other negatively correlated terms. This likely reflects the influence of strong genetic effects, such as those at the *MKT1/SAL1* locus described above but also the effect of the *HAP1* hotspot (Brem *et al*., 2002) on mitochondrial genes in the BYxRM collection. While the different parental alleles at the *MKT1/SAL1* locus led to a highly significant change in the PT score that was only absent for proteins in a limited number of functional modules (see main text), allelic differences at the *HAP1* hotspot did not display any directional effect on the PT score, despite affecting proteins annotated for mitochondrial functions in a similar manner as *MKT1/SAL1* alleles. We propose that a distinction between primary and secondary effect would be most suited to explain this apparent contradiction. Specifically, a primary effect on mitochondrial function may or may not lead to a reorganization of the cellular configuration in terms of the PT network state, dependent on the timing, reach and character of the perturbation. In this context, it is also of importance to consider the time of the measurement, at which the state of the cellular network is observed, which certainly does not exclude an effect at other time-points during the growth of a yeast culture (such as close to or during diauxic shift or even during subsequent stationary phase growth).

For example, we noted that transcription by RNA polymerase I and III but not II were among the positively correlated terms. Similar correlations have been reported recently for fission yeast cells grown on different carbon sources (Kleijn et al., 2021) and RNA pol III transcription seems to be required for glycolysis indicating a potential feedback loop with PKA or TOR signaling (Szatkowska et al., 2019). The strong negative correlation with the term “oligosaccharide metabolic process” included low levels of trehalose synthetase subunits in strains with high PT score, as expected. However, we observed that isoforms of phosphoglucomutase differed in this regard: While Pgm2p was reduced with high scores, as expected (Howard et al., further REFs), Pgm1p was significantly increased (LM beta = 0.49, q < 1E-14), hinting to an unknown role of this enzyme during growth on glucose. We also noted overall slightly positive correlation of the PT score with proteins annotated for transposition. Among these, RPC40, an RNA polymerase III subunit known for its role in transposon insertion (Bridier-Nahmias, 2015), was strongly associated with high PT scores (LM beta = +0.56, q < 1E-18). Moreover, we found strong predictive power, but variable directionality of correlation for subsets of proteins annotated for protein phosphorylation and dephosphorylation. These correlations may yield valuable insight into the regulation of kinases and phosphatase subunits.

The strong coverage of phosphorylation events across segregants in our dataset and interpretation in light of the PT score provided a wealth of information about many regulatory processes. As an example, we further scrutinized strong correlations between the PT score and phosphorylation events in proteins that are annotated for functions in histone modification. We noticed that several proteins involved in deposition, reading and removal of histone H3 lysine 36 (H3K36) methylation contained phosphopeptides that were strongly predicted by the PT score (Fig. 2H or suppl. Fig). It has been reported that H3K36 trimethylation is important to prevent bi-directional (cryptic) transcription within gene bodies. In turn, the expression of nutrient-sensitive as well as stress response genes depends on the presence of the histone methylase Set2 (McDaniel, 2017; Separovich, 2022). Set2 mediates co-transcriptional histone modification, which was further shown to depend on the cyclin-dependent kinase Bur1 and the Bur2 cyclin (Hossain, 2013). The process initiated by Set2p further involves recruitment of the Rpd3S histone deacetylase complex (Reim, 2020). Set2 interacts with the histone chaperone Spt6 and the Iws1/Spn1 component of the RNA polymerase II elongation complex and this interaction is crucial for the expression of highly transcribed mRNAs such as those encoding ribosomal proteins (Reim, 2020). Conversely, demethylases Jhd1p, Jhd2p, Rph1p, and Gis1p remove H3K36 marks (Separovich, 2022). We found strong correlations between the PT score and phosphopeptides in the BUR kinase subunit Sgv1, Bur2, Set2, Iws1/Spn1 and its binding partner Spt6, the Sin3 component of the Rpd3S complex, Rph1 as well as Gis1. Furthermore, we found strong correlations with phosphopeptides in Chd1, a subunit of the SAGA complex, which recognizes H3K4 methylation via its chromodomain (Pray-Grant, 2005). Chd1 was also shown to bind to H3K36 methylated sites (Smolle, 2012) and loss of Chd1 alone or in combination with loss of Iws1/Spn1 resulted in cryptic transcription (Quan, 2010). Our observation that Chd1 phosphorylation strongly correlated with PT score variability supports recent reports that SAGA complex recruitment may be regulated in a TORC2-dependent manner (Cohen, 2022) and provides another intriguing link to transcriptional fidelity.

## Supplemental figures

**Figure S1.**
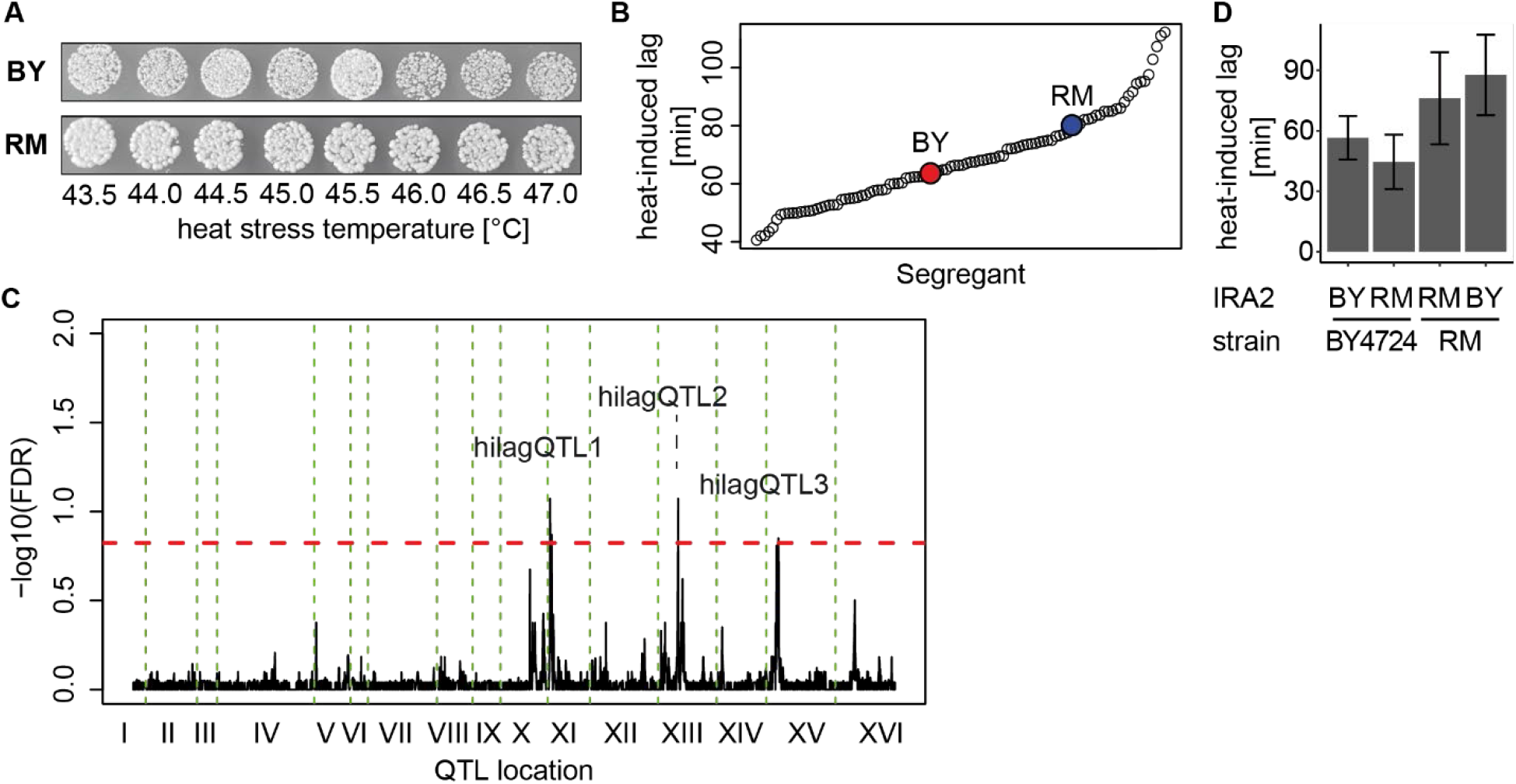
**A** Viability test for heat stress treatment. Samples from exponentially growing cultures of BY or RM were subjected to transient heat stress treatment (ramping from 25 °C to indicated temperature at 1 K/s, followed by 8 min exposure at constant peak temperature) and spotted onto YPD plates following appropriate dilution. Spots were photographed after 1.5 days at 30 °C. **B** Distribution of heat-induced lag measurements across 100 segregants and the parental strains as indicated. **C** QTL mapping result for heat-induced lag. Dashed red line represents 15 % FDR threshold based on comparison between mapping of true against permuted trait values. Loci that passed the threshold are indicated. **D** Heat-induced lag measurements (n = 4) in BY4724, which is closely related to the parental strain BY4716, and in RM11-1a as well as in derivatives after allele-swapping of *IRA2* (Smith and Kruglyak, 2008).

**Figure S2.**
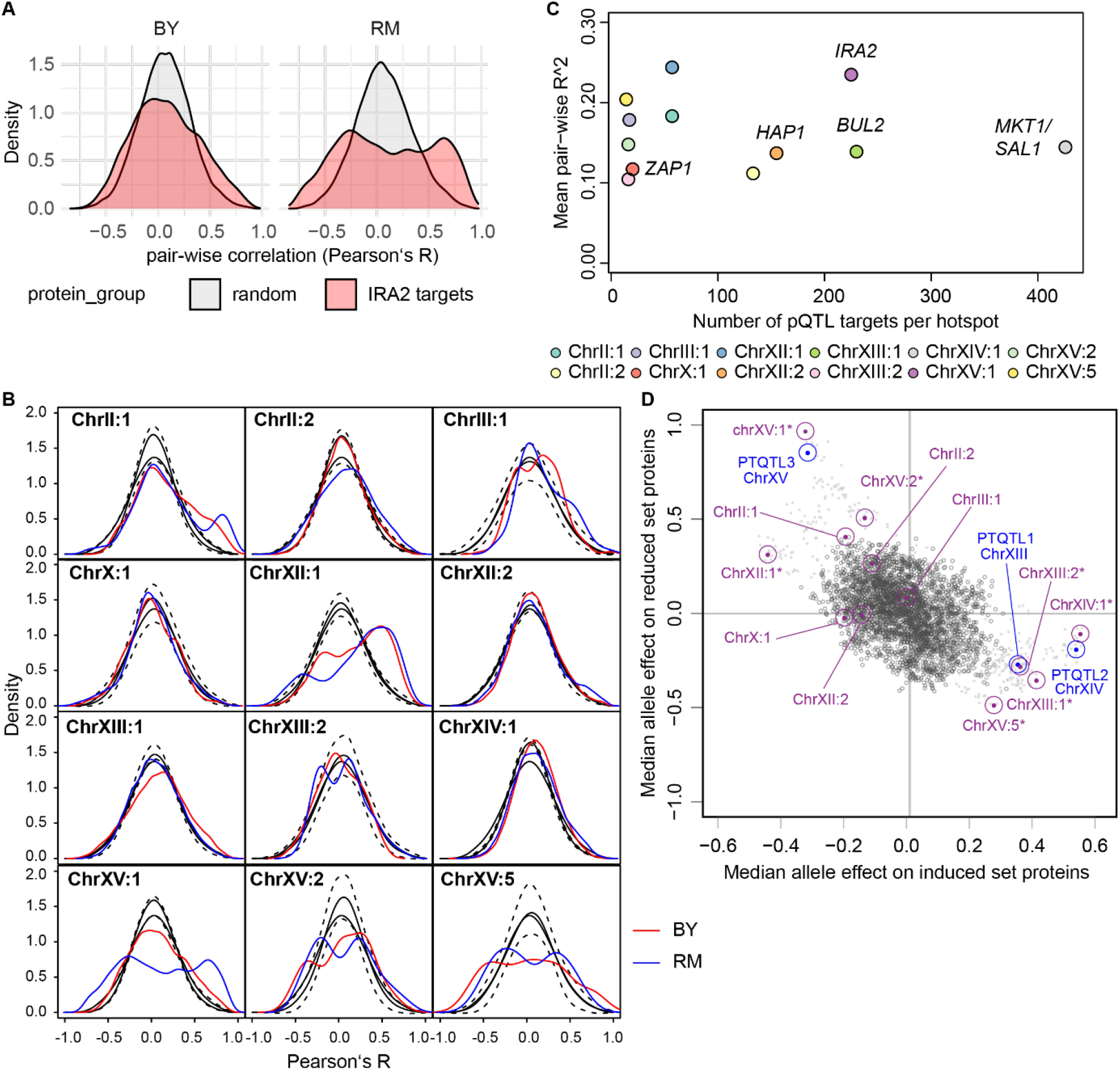
**A** Test for coordinated expression among protein abundance targets of the *IRA2* pQTL hotspot. Pair-wise correlation among 225 proteins shown separately for sets of strains carrying either the BY or the RM allele at the marker corresponding to the *IRA2* gene. The grey colored area represents a representative example of the distribution of pair-wise correlations in abundance-matched but otherwise random sets of proteins. The average distribution as well as standard deviation of the frequency distribution (200 bins) are shown in (B). **B** Same test for coordination as in (A) but for each of 12 pQTL hotspots as described in (Grossbach *et al*., 2022). Pair-wise correlations between protein abundance targets of the indicated hotspot were calculated in sets of strains split by the corresponding most significant marker (BY: red curve, RM: blue curve). Black solid lines show the average distribution of pair-wise correlations in random sets of proteins that were matched in abundance to the targets of the respective hotspot. These pair-wise correlations were calculated in the same sets of strains as used for the actual targets, represented by two different solid lines. Dashed lines show the maximum and minimum among 1000 random samples of proteins, calculated for 200 bins across the range from −1 to +1. **C** Comparison of coordinated expression of pQTL hotspot targets (average of all pair-wise correlations as shown in (B)) and the number of targets for each hotspot. **D** Comparison of median allelic effect at each of 3,593 genetic markers on the abundance of induced and repressed PT marker proteins. Significant QTL for the PT score are marked by blue circles and pQTL hotspots are marked by purple circles (asterisk indicates significant effect on the PT score at q < 0.05). Small dots are markers that are in linkage disequilibrium with a lead marker of any of the hotspots with significant PT score differences (R > 0.35).

**Figure S3.**
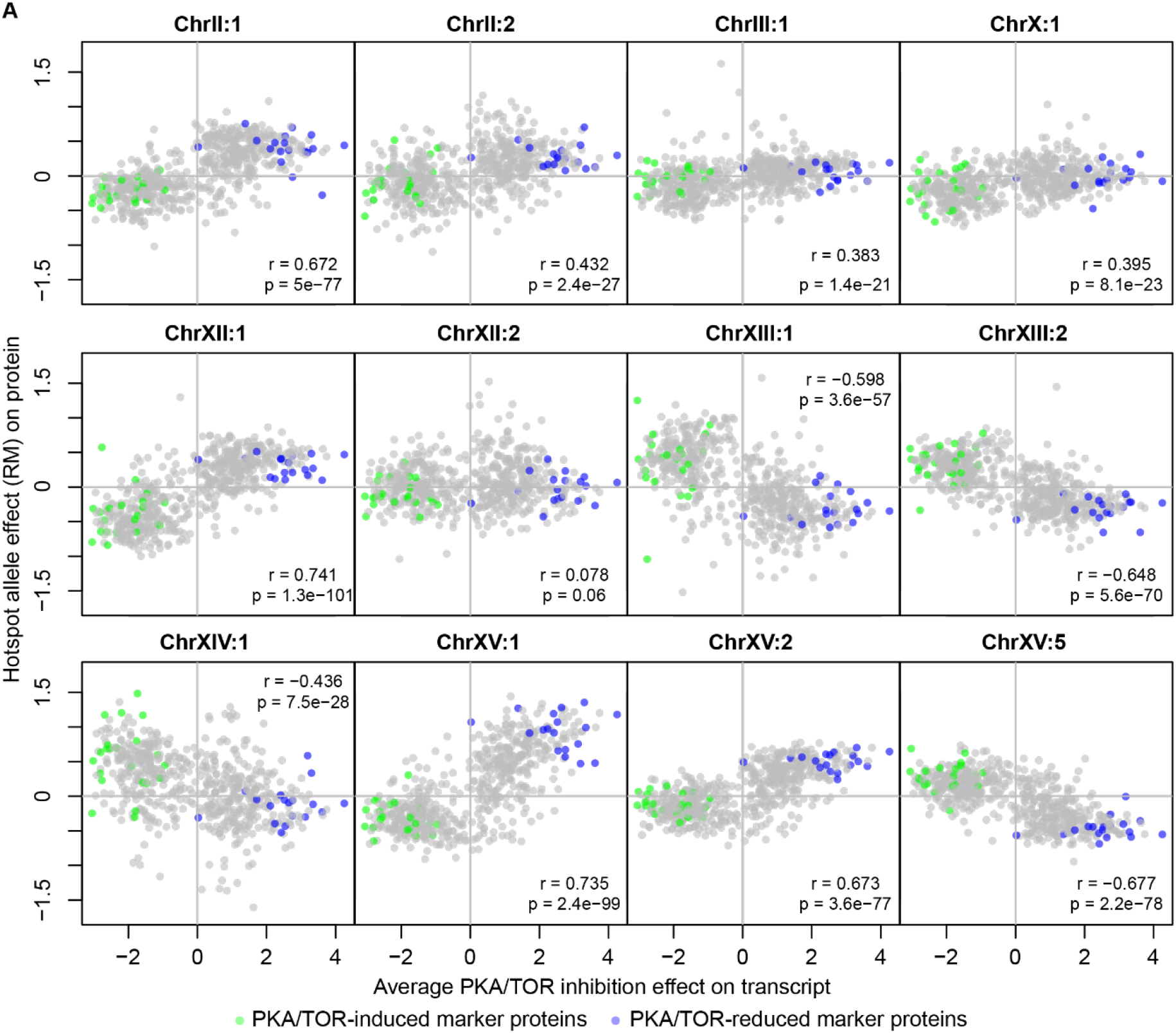
**A** Comparison of protein abundance changes associated with allele differences at 12 pQTL hotspots to the effect of chemical inhibition of PKA and TOR signaling pathways on transcript abundance (Kunkel *et al*., 2019). The effect of chemical inhibition as shown here represents the average effect following 20 min inhibition of either pathway. Marker genes used for calculation of the PT score are highlighted.

**Figure S4.**
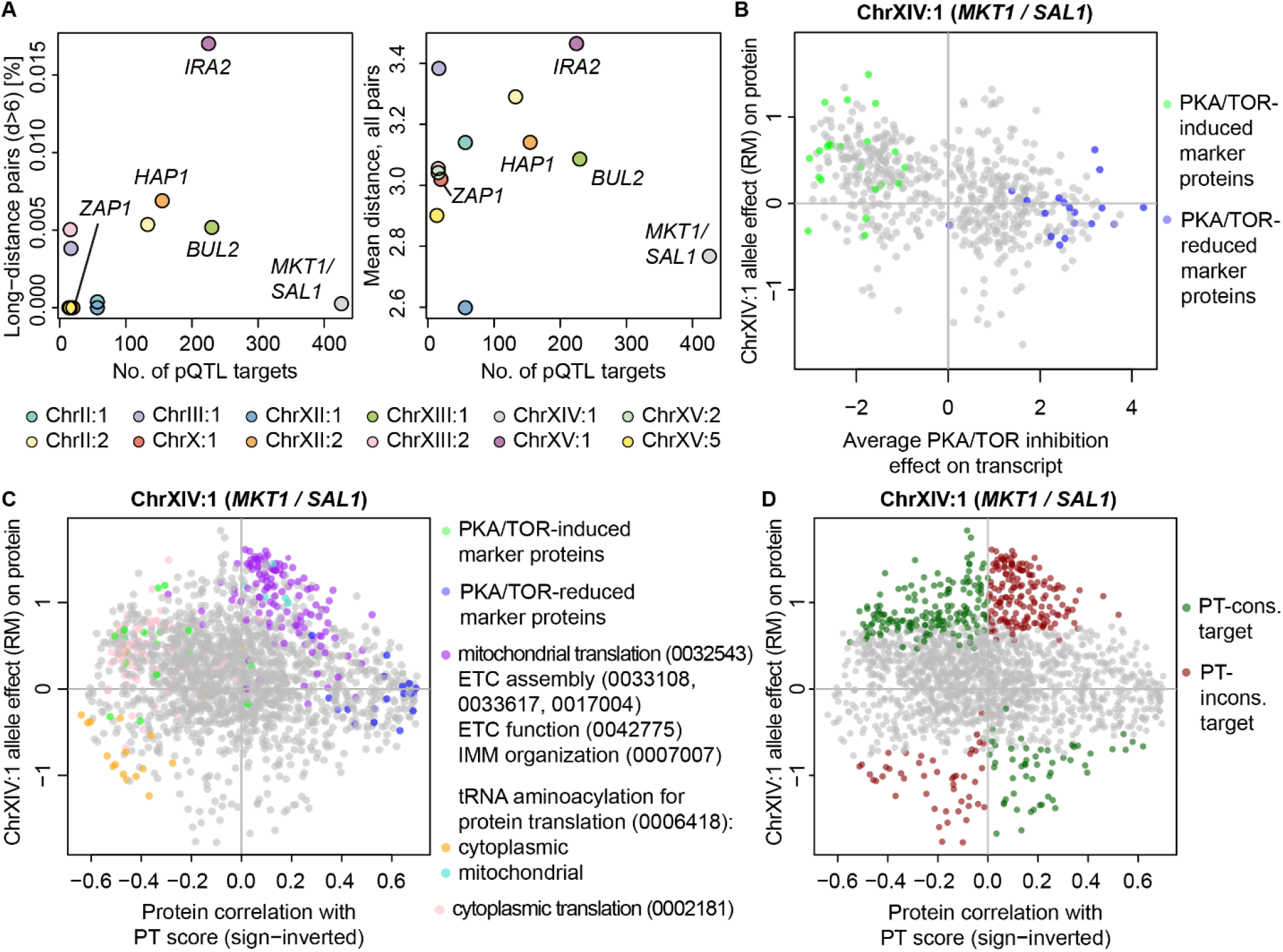
**A** Pair-wise correlation among targets of pQTL hotspot “ChrX:I” (*ZAP1*) after splitting by the direction of abundance change in sets of strains split by the corresponding most significant genetic marker. **B** Comparison of shortest path distances as in Fig. 4A for all pair-wise combinations of targets of the indicated pQTL hotspots to the number of targets. Left panel: Fraction of long-distance pairs with a distance of more than 6 steps. Right panel: Mean distance across all pairs. **C** Analysis of protein abundance changes due to allelic differences at hotspot ChrXIV:1, which spans the *MKT1* and *SAL1* loci, by partitioning of proteins into GO terms as indicated. Comparison to chemical inhibition as in Fig. S3A (left panel) and comparison of allele-dependent abundance changes to global correlation of respective protein abundances with the PT score across BYxRM segregants (right panel). The x-axis shows the beta coefficient of the PT score as a predictor of protein abundance in a combined linear model with the genotype at the hotspot ChrXIV:1 locus to correct for the effect of the locus itself.

**Figure S5.**
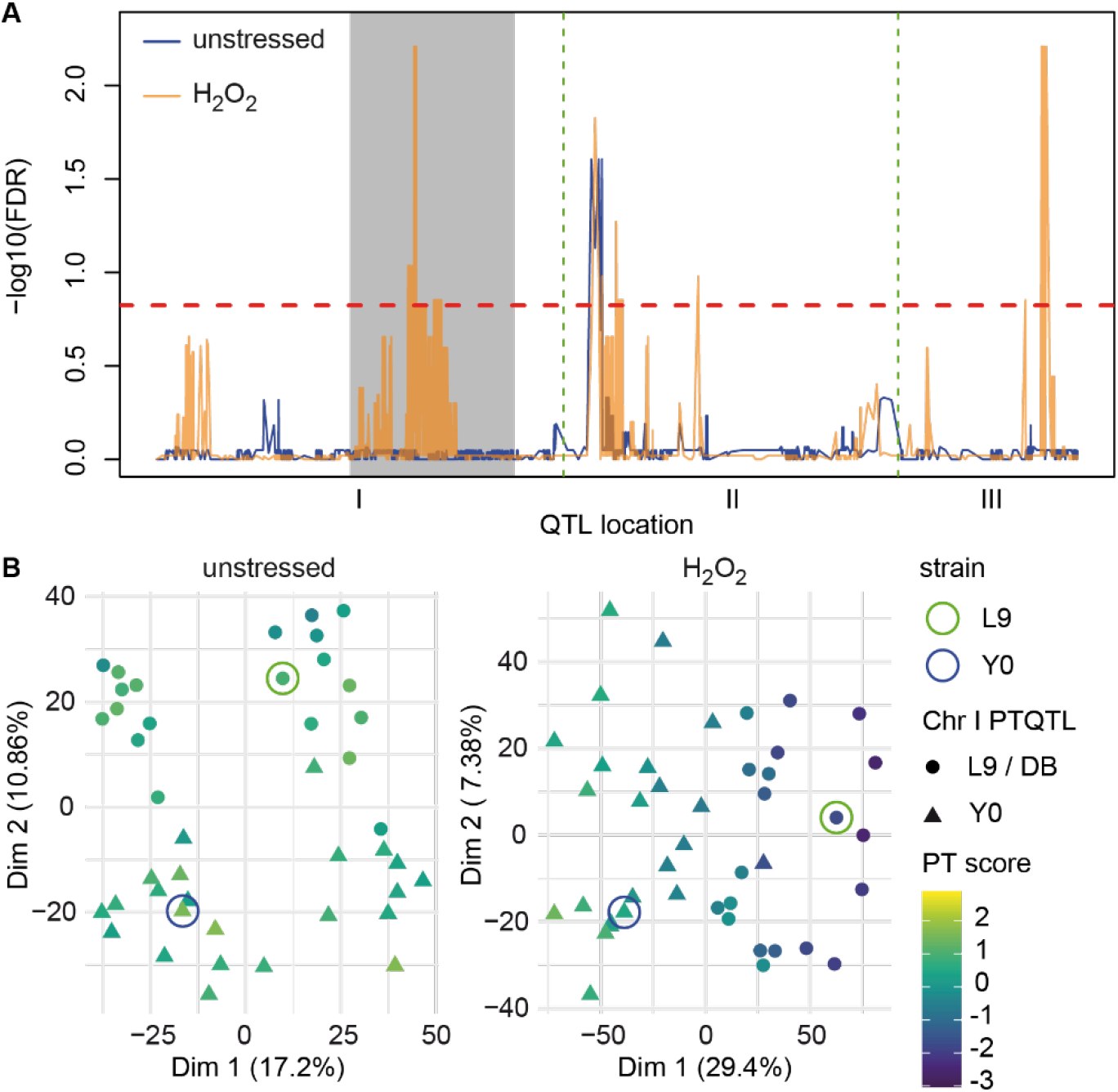
**A** QTL mapping of the PT score in the fission yeast 3-way cross in two conditions, as indicated. 15 % FDR indicated by red dashed horizontal line. Shaded area represents inverted region of Chromosome I in the Y0 parental strain. **B** PCA based on transcriptome variability for 43 segregants between the L9 and Y0 parental strains (cross R1, parental strains highlighted) in two conditions. Strains are colored by PT score and shape represents allele identity at the most significant marker of PTQTL1 for strains in H_2_O_2_.

## References

Abe, F., and Iida, H. (2003). Pressure-induced differential regulation of the two tryptophan permeases Tat1 and Tat2 by ubiquitin ligase Rsp5 and its binding proteins, Bul1 and Bul2. Mol Cell Biol 23, 7566–7584. 10.1128/MCB.23.21.7566-7584.2003.

Abrams, M.B., Chuong, J.N., AlZaben, F., Dubin, C.A., Skerker, J.M., and Brem, R.B. (2022). Barcoded reciprocal hemizygosity analysis via sequencing illuminates the complex genetic basis of yeast thermotolerance. G3 (Bethesda) 12. 10.1093/g3journal/jkab412.

Balakrishnan, R., de Silva, R.T., Hwa, T., and Cremer, J. (2021). Suboptimal resource allocation in changing environments constrains response and growth in bacteria. Mol Syst Biol 17, e10597. 10.15252/msb.202110597.

Basan, M., Hui, S., Okano, H., Zhang, Z., Shen, Y., Williamson, J.R., and Hwa, T. (2015). Overflow metabolism in Escherichia coli results from efficient proteome allocation. Nature 528, 99–104. 10.1038/nature15765.

Belenky, P.A., Moga, T.G., and Brenner, C. (2008). Saccharomyces cerevisiae YOR071C encodes the high affinity nicotinamide riboside transporter Nrt1. J Biol Chem 283, 8075–8079. 10.1074/jbc.C800021200.

Bhattacharya, D., Azambuja, A.P., and Simoes-Costa, M. (2020). Metabolic Reprogramming Promotes Neural Crest Migration via Yap/Tead Signaling. Dev Cell 53, 199–211 e196. 10.1016/j.devcel.2020.03.005.

Bolger, A.M., Lohse, M., and Usadel, B. (2014). Trimmomatic: a flexible trimmer for Illumina sequence data. Bioinformatics 30, 2114–2120. 10.1093/bioinformatics/btu170.

Boyle, E.A., Li, Y.I., and Pritchard, J.K. (2017). An Expanded View of Complex Traits: From Polygenic to Omnigenic. Cell 169, 1177–1186. 10.1016/j.cell.2017.05.038.

Brauer, M.J., Saldanha, A.J., Dolinski, K., and Botstein, D. (2005). Homeostatic adjustment and metabolic remodeling in glucose-limited yeast cultures. Mol Biol Cell 16, 2503–2517. 10.1091/mbc.e04-11-0968.

Brem, R.B., Yvert, G., Clinton, R., and Kruglyak, L. (2002). Genetic dissection of transcriptional regulation in budding yeast. Science 296, 752–755. 10.1126/science.1069516.

Broach, J.R. (2012). Nutritional control of growth and development in yeast. Genetics 192, 73–105. 10.1534/genetics.111.135731.

Campbell, K., Westholm, J., Kasvandik, S., Di Bartolomeo, F., Mormino, M., and Nielsen, J. (2020). Building blocks are synthesized on demand during the yeast cell cycle. Proc Natl Acad Sci U S A 117, 7575–7583. 10.1073/pnas.1919535117.

Caspeta, L., Chen, Y., and Nielsen, J. (2016). Thermotolerant yeasts selected by adaptive evolution express heat stress response at 30 degrees C. Sci Rep 6, 27003. 10.1038/srep27003.

Chen, J.C., and Powers, T. (2006). Coordinate regulation of multiple and distinct biosynthetic pathways by TOR and PKA kinases in S. cerevisiae. Curr Genet 49, 281–293. 10.1007/s00294-005-0055-9.

Chen, X., Wang, G., Zhang, Y., Dayhoff-Brannigan, M., Diny, N.L., Zhao, M., He, G., Sing, C.N., Metz, K.A., Stolp, Z.D., et al. (2018). Whi2 is a conserved negative regulator of TORC1 in response to low amino acids. PLoS Genet 14, e1007592. 10.1371/journal.pgen.1007592.

Chen, Y., Zhu, J., Lum, P.Y., Yang, X., Pinto, S., MacNeil, D.J., Zhang, C., Lamb, J., Edwards, S., Sieberts, S.K., et al. (2008). Variations in DNA elucidate molecular networks that cause disease. Nature 452, 429–435. 10.1038/nature06757.

Cherkasov, V., Hofmann, S., Druffel-Augustin, S., Mogk, A., Tyedmers, J., Stoecklin, G., and Bukau, B. (2013). Coordination of translational control and protein homeostasis during severe heat stress. Curr Biol 23, 2452–2462. 10.1016/j.cub.2013.09.058.

Cherry, J.M., Hong, E.L., Amundsen, C., Balakrishnan, R., Binkley, G., Chan, E.T., Christie, K.R., Costanzo, M.C., Dwight, S.S., Engel, S.R., et al. (2012). Saccharomyces Genome Database: the genomics resource of budding yeast. Nucleic Acids Res 40, D700–705. 10.1093/nar/gkr1029.

Clement-Ziza, M., Marsellach, F.X., Codlin, S., Papadakis, M.A., Reinhardt, S., Rodriguez-Lopez, M., Martin, S., Marguerat, S., Schmidt, A., Lee, E., et al. (2014). Natural genetic variation impacts expression levels of coding, non-coding, and antisense transcripts in fission yeast. Mol Syst Biol 10, 764. 10.15252/msb.20145123.

Conrad, M., Schothorst, J., Kankipati, H.N., Van Zeebroeck, G., Rubio-Texeira, M., and Thevelein, J.M. (2014). Nutrient sensing and signaling in the yeast Saccharomyces cerevisiae. FEMS Microbiol Rev 38, 254–299. 10.1111/1574-6976.12065.

Davidson, J.F., Whyte, B., Bissinger, P.H., and Schiestl, R.H. (1996). Oxidative stress is involved in heat-induced cell death in Saccharomyces cerevisiae. Proc Natl Acad Sci U S A 93, 5116–5121. 10.1073/pnas.93.10.5116.

Dilova, I., Aronova, S., Chen, J.C., and Powers, T. (2004). Tor signaling and nutrient-based signals converge on Mks1p phosphorylation to regulate expression of Rtg1.Rtg3p-dependent target genes. J Biol Chem 279, 46527–46535. 10.1074/jbc.M409012200.

Dimitrov, L.N., Brem, R.B., Kruglyak, L., and Gottschling, D.E. (2009). Polymorphisms in multiple genes contribute to the spontaneous mitochondrial genome instability of Saccharomyces cerevisiae S288C strains. Genetics 183, 365–383. 10.1534/genetics.109.104497.

Duran, R.V., Oppliger, W., Robitaille, A.M., Heiserich, L., Skendaj, R., Gottlieb, E., and Hall, M.N. (2012). Glutaminolysis activates Rag-mTORC1 signaling. Mol Cell 47, 349–358. 10.1016/j.molcel.2012.05.043.

Elbing, K., Larsson, C., Bill, R.M., Albers, E., Snoep, J.L., Boles, E., Hohmann, S., and Gustafsson, L. (2004). Role of hexose transport in control of glycolytic flux in Saccharomyces cerevisiae. Appl Environ Microbiol 70, 5323–5330. 10.1128/AEM.70.9.5323-5330.2004.

Ellegren, H. (2005). Evolution: natural selection in the evolution of humans and chimps. Curr Biol 15, R919–922. 10.1016/j.cub.2005.10.060.

Franzmann, T.M., and Alberti, S. (2019). Protein Phase Separation as a Stress Survival Strategy. Cold Spring Harb Perspect Biol 11. 10.1101/cshperspect.a034058.

Frei, T., Cella, F., Tedeschi, F., Gutierrez, J., Stan, G.B., Khammash, M., and Siciliano, V. (2020). Characterization and mitigation of gene expression burden in mammalian cells. Nat Commun 11, 4641. 10.1038/s41467-020-18392-x.

Gasch, A.P., Spellman, P.T., Kao, C.M., Carmel-Harel, O., Eisen, M.B., Storz, G., Botstein, D., and Brown, P.O. (2000). Genomic expression programs in the response of yeast cells to environmental changes. Mol Biol Cell 11, 4241–4257. 10.1091/mbc.11.12.4241.

Gibney, P.A., Lu, C., Caudy, A.A., Hess, D.C., and Botstein, D. (2013). Yeast metabolic and signaling genes are required for heat-shock survival and have little overlap with the heat-induced genes. Proc Natl Acad Sci U S A 110, E4393–4402. 10.1073/pnas.1318100110.

Gonzalez, A., and Hall, M.N. (2017). Nutrient sensing and TOR signaling in yeast and mammals. EMBO J 36, 397–408. 10.15252/embj.201696010.

Gonzalez, A., Hall, M.N., Lin, S.C., and Hardie, D.G. (2020). AMPK and TOR: The Yin and Yang of Cellular Nutrient Sensing and Growth Control. Cell Metab 31, 472–492. 10.1016/j.cmet.2020.01.015.

Grossbach, J., Gillet, L., Clement-Ziza, M., Schmalohr, C.L., Schubert, O.T., Schutter, M., Mawer, J.S.P., Barnes, C.A., Bludau, I., Weith, M., et al. (2022). The impact of genomic variation on protein phosphorylation states and regulatory networks. Mol Syst Biol 18, e10712. 10.15252/msb.202110712.

Han, J.D. (2008). Understanding biological functions through molecular networks. Cell Res 18, 224–237. 10.1038/cr.2008.16.

Hara, K., Yonezawa, K., Weng, Q.P., Kozlowski, M.T., Belham, C., and Avruch, J. (1998). Amino acid sufficiency and mTOR regulate p70 S6 kinase and eIF-4E BP1 through a common effector mechanism. J Biol Chem 273, 14484–14494. 10.1074/jbc.273.23.14484.

Hu, X.P., Dourado, H., Schubert, P., and Lercher, M.J. (2020). The protein translation machinery is expressed for maximal efficiency in Escherichia coli. Nat Commun 11, 5260. 10.1038/s41467-020-18948-x.

Jarolim, S., Ayer, A., Pillay, B., Gee, A.C., Phrakaysone, A., Perrone, G.G., Breitenbach, M., and Dawes, I.W. (2013). Saccharomyces cerevisiae genes involved in survival of heat shock. G3 (Bethesda) 3, 2321–2333. 10.1534/g3.113.007971.

Kahm, M., Hasenbrink, G., Lichtenberg-Fraté, H., Ludwig, J., and Kschischo, M. (2010). grofit: Fitting Biological Growth Curves withR. Journal of Statistical Software 33. 10.18637/jss.v033.i07.

Kamrad, S., Grossbach, J., Rodriguez-Lopez, M., Mulleder, M., Townsend, S., Cappelletti, V., Stojanovski, G., Correia-Melo, C., Picotti, P., Beyer, A., et al. (2020). Pyruvate kinase variant of fission yeast tunes carbon metabolism, cell regulation, growth and stress resistance. Mol Syst Biol 16, e9270. 10.15252/msb.20199270.

Kaya, A., Phua, C.Z.J., Lee, M., Wang, L., Tyshkovskiy, A., Ma, S., Barre, B., Liu, W., Harrison, B.R., Zhao, X., et al. (2021). Evolution of natural lifespan variation and molecular strategies of extended lifespan in yeast. Elife 10. 10.7554/eLife.64860.

Kingsbury, J.M., Sen, N.D., and Cardenas, M.E. (2015). Branched-Chain Aminotransferases Control TORC1 Signaling in Saccharomyces cerevisiae. PLoS Genet 11, e1005714. 10.1371/journal.pgen.1005714.

Kleijn, I.T., Martinez-Segura, A., Bertaux, F., Saint, M., Kramer, H., Shahrezaei, V., and Marguerat, S. (2022). Growth-rate-dependent and nutrient-specific gene expression resource allocation in fission yeast. Life Sci Alliance 5. 10.26508/lsa.202101223.

Kunkel, J., Luo, X., and Capaldi, A.P. (2019). Integrated TORC1 and PKA signaling control the temporal activation of glucose-induced gene expression in yeast. Nat Commun 10, 3558. 10.1038/s41467-019-11540-y.

Kwan, E.X., Foss, E., Kruglyak, L., and Bedalov, A. (2011). Natural polymorphism in BUL2 links cellular amino acid availability with chronological aging and telomere maintenance in yeast. PLoS Genet 7, e1002250. 10.1371/journal.pgen.1002250.

Langmead, B., and Salzberg, S.L. (2012). Fast gapped-read alignment with Bowtie 2. Nat Methods 9, 357–359. 10.1038/nmeth.1923.

Leadsham, J.E., Miller, K., Ayscough, K.R., Colombo, S., Martegani, E., Sudbery, P., and Gourlay, C.W. (2009). Whi2p links nutritional sensing to actin-dependent Ras-cAMP-PKA regulation and apoptosis in yeast. J Cell Sci 122, 706–715. 10.1242/jcs.042424.

Lee, P., Kim, M.S., Paik, S.-M., Choi, S.-H., Cho, B.-R., and Hahn, J.-S. (2013). Rim15-dependent activation of Hsf1 and Msn2/4 transcription factors by direct phosphorylation inSaccharomyces cerevisiae. FEBS Letters 587, 3648–3655. 10.1016/j.febslet.2013.10.004.

Liberti, M.V., and Locasale, J.W. (2016). The Warburg Effect: How Does it Benefit Cancer Cells? Trends Biochem Sci 41, 211–218. 10.1016/j.tibs.2015.12.001.

Lippman, S.I., and Broach, J.R. (2009). Protein kinase A and TORC1 activate genes for ribosomal biogenesis by inactivating repressors encoded by Dot6 and its homolog Tod6. Proc Natl Acad Sci U S A 106, 19928–19933. 10.1073/pnas.0907027106.

Liu, H., Styles, C.A., and Fink, G.R. (1996). Saccharomyces cerevisiae S288C has a mutation in FLO8, a gene required for filamentous growth. Genetics 144, 967–978. 10.1093/genetics/144.3.967.

Liu, X., Li, Y.I., and Pritchard, J.K. (2019). Trans Effects on Gene Expression Can Drive Omnigenic Inheritance. Cell 177, 1022–1034 e1026. 10.1016/j.cell.2019.04.014.

Love, M.I., Huber, W., and Anders, S. (2014). Moderated estimation of fold change and dispersion for RNA-seq data with DESeq2. Genome Biol 15, 550. 10.1186/s13059-014-0550-8.

Lu, J.M., Deschenes, R.J., and Fassler, J.S. (2004). Role for the Ran binding protein, Mog1p, in Saccharomyces cerevisiae SLN1-SKN7 signal transduction. Eukaryot Cell 3, 1544–1556. 10.1128/EC.3.6.1544-1556.2004.

Lyne, R., Burns, G., Mata, J., Penkett, C.J., Rustici, G., Chen, D., Langford, C., Vetrie, D., and Bahler, J. (2003). Whole-genome microarrays of fission yeast: characteristics, accuracy, reproducibility, and processing of array data. BMC Genomics 4, 27. 10.1186/1471-2164-4-27.

Lyons, T.J., Gasch, A.P., Gaither, L.A., Botstein, D., Brown, P.O., and Eide, D.J. (2000). Genome-wide characterization of the Zap1p zinc-responsive regulon in yeast. Proc Natl Acad Sci U S A 97, 7957–7962. 10.1073/pnas.97.14.7957.

Malecki, M., Kamrad, S., Ralser, M., and Bahler, J. (2020). Mitochondrial respiration is required to provide amino acids during fermentative proliferation of fission yeast. EMBO Rep 21, e50845. 10.15252/embr.202050845.

McDaniel, S.L., and Strahl, B.D. (2017). Shaping the cellular landscape with Set2/SETD2 methylation. Cell Mol Life Sci 74, 3317–3334. 10.1007/s00018-017-2517-x.

Merhi, A., and Andre, B. (2012). Internal amino acids promote Gap1 permease ubiquitylation via TORC1/Npr1/14-3-3-dependent control of the Bul arrestin-like adaptors. Mol Cell Biol 32, 4510–4522. 10.1128/MCB.00463-12.

Michaelson, J.J., Alberts, R., Schughart, K., and Beyer, A. (2010). Data-driven assessment of eQTL mapping methods. BMC Genomics 11, 502. 10.1186/1471-2164-11-502.

Molenaar, D., van Berlo, R., de Ridder, D., and Teusink, B. (2009). Shifts in growth strategies reflect tradeoffs in cellular economics. Mol Syst Biol 5, 323. 10.1038/msb.2009.82.

Mori, M., Zhang, Z., Banaei-Esfahani, A., Lalanne, J.B., Okano, H., Collins, B.C., Schmidt, A., Schubert, O.T., Lee, D.S., Li, G.W., et al. (2021). From coarse to fine: the absolute Escherichia coli proteome under diverse growth conditions. Mol Syst Biol 17, e9536. 10.15252/msb.20209536.

Nguyen Ba, A.N., Lawrence, K.R., Rego-Costa, A., Gopalakrishnan, S., Temko, D., Michor, F., and Desai, M.M. (2022). Barcoded bulk QTL mapping reveals highly polygenic and epistatic architecture of complex traits in yeast. Elife 11. 10.7554/eLife.73983.

Oliveira, A.P., Dimopoulos, S., Busetto, A.G., Christen, S., Dechant, R., Falter, L., Haghir Chehreghani, M., Jozefczuk, S., Ludwig, C., Rudroff, F., et al. (2015). Inferring causal metabolic signals that regulate the dynamic TORC1-dependent transcriptome. Mol Syst Biol 11, 802. 10.15252/msb.20145475.

Ottens, F., Franz, A., and Hoppe, T. (2021). Build-UPS and break-downs: metabolism impacts on proteostasis and aging. Cell Death Differ 28, 505–521. 10.1038/s41418-020-00682-y.

Paulo, J.A., O’Connell, J.D., Everley, R.A., O’Brien, J., Gygi, M.A., and Gygi, S.P. (2016). Quantitative mass spectrometry-based multiplexing compares the abundance of 5000 S. cerevisiae proteins across 10 carbon sources. J Proteomics 148, 85–93. 10.1016/j.jprot.2016.07.005.

Pedruzzi, I., Dubouloz, F., Cameroni, E., Wanke, V., Roosen, J., Winderickx, J., and De Virgilio, C. (2003). TOR and PKA Signaling Pathways Converge on the Protein Kinase Rim15 to Control Entry into G0. Molecular Cell 12, 1607–1613. 10.1016/s1097-2765(03)00485-4.

Peeters, K., Van Leemputte, F., Fischer, B., Bonini, B.M., Quezada, H., Tsytlonok, M., Haesen, D., Vanthienen, W., Bernardes, N., Gonzalez-Blas, C.B., et al. (2017). Fructose-1,6-bisphosphate couples glycolytic flux to activation of Ras. Nat Commun 8, 922. 10.1038/s41467-017-01019-z.

Peters, L.A., Perrigoue, J., Mortha, A., Iuga, A., Song, W.M., Neiman, E.M., Llewellyn, S.R., Di Narzo, A., Kidd, B.A., Telesco, S.E., et al. (2017). A functional genomics predictive network model identifies regulators of inflammatory bowel disease. Nat Genet 49, 1437–1449. 10.1038/ng.3947.

Plank, M. (2022). Interaction of TOR and PKA Signaling in S. cerevisiae. Biomolecules 12, 210. 10.3390/biom12020210.

Plank, M., Perepelkina, M., Muller, M., Vaga, S., Zou, X., Bourgoint, C., Berti, M., Saarbach, J., Haesendonckx, S., Winssinger, N., et al. (2020). Chemical Genetics of AGC-kinases Reveals Shared Targets of Ypk1, Protein Kinase A and Sch9. Mol Cell Proteomics 19, 655–671. 10.1074/mcp.RA120.001955.

Quinlan, A.R., and Hall, I.M. (2010). BEDTools: a flexible suite of utilities for comparing genomic features. Bioinformatics 26, 841–842. 10.1093/bioinformatics/btq033.

Ramachandran, V., and Herman, P.K. (2011). Antagonistic interactions between the cAMP-dependent protein kinase and Tor signaling pathways modulate cell growth in Saccharomyces cerevisiae. Genetics 187, 441–454. 10.1534/genetics.110.123372.

Reinders, A., Burckert, N., Boller, T., Wiemken, A., and De Virgilio, C. (1998). Saccharomyces cerevisiae cAMP-dependent protein kinase controls entry into stationary phase through the Rim15p protein kinase. Genes Dev 12, 2943–2955. 10.1101/gad.12.18.2943.

Rhind, N., Chen, Z., Yassour, M., Thompson, D.A., Haas, B.J., Habib, N., Wapinski, I., Roy, S., Lin, M.F., Heiman, D.I., et al. (2011). Comparative functional genomics of the fission yeasts. Science 332, 930–936. 10.1126/science.1203357.

Rodriguez-Lopez, M., Cotobal, C., Fernandez-Sanchez, O., Borbaran Bravo, N., Oktriani, R., Abendroth, H., Uka, D., Hoti, M., Wang, J., Zaratiegui, M., and Bahler, J. (2016). A CRISPR/Cas9-based method and primer design tool for seamless genome editing in fission yeast. Wellcome Open Res 1, 19. 10.12688/wellcomeopenres.10038.3.

Schadt, E.E. (2009). Molecular networks as sensors and drivers of common human diseases. Nature 461, 218–223. 10.1038/nature08454.

Separovich, R.J., Wong, M.W.M., Bartolec, T.K., Hamey, J.J., and Wilkins, M.R. (2022). Site-specific Phosphorylation of Histone H3K36 Methyltransferase Set2p and Demethylase Jhd1p is Required for Stress Responses in Saccharomyces cerevisiae. J Mol Biol 434, 167500. 10.1016/j.jmb.2022.167500.

Shimobayashi, M., and Hall, M.N. (2016). Multiple amino acid sensing inputs to mTORC1. Cell Res 26, 7–20. 10.1038/cr.2015.146.

Sinha, H., David, L., Pascon, R.C., Clauder-Munster, S., Krishnakumar, S., Nguyen, M., Shi, G., Dean, J., Davis, R.W., Oefner, P.J., et al. (2008). Sequential elimination of major-effect contributors identifies additional quantitative trait loci conditioning high-temperature growth in yeast. Genetics 180, 1661–1670. 10.1534/genetics.108.092932.

Sinha, H., Nicholson, B.P., Steinmetz, L.M., and McCusker, J.H. (2006). Complex genetic interactions in a quantitative trait locus. PLoS Genet 2, e13. 10.1371/journal.pgen.0020013.

Smith, E.N., and Kruglyak, L. (2008). Gene-environment interaction in yeast gene expression. PLoS Biol 6, e83. 10.1371/journal.pbio.0060083.

Soulard, A., Cremonesi, A., Moes, S., Schutz, F., Jeno, P., and Hall, M.N. (2010). The rapamycin-sensitive phosphoproteome reveals that TOR controls protein kinase A toward some but not all substrates. Mol Biol Cell 21, 3475–3486. 10.1091/mbc.E10-03-0182.

Steinmetz, L.M., Sinha, H., Richards, D.R., Spiegelman, J.I., Oefner, P.J., McCusker, J.H., and Davis, R.W. (2002). Dissecting the architecture of a quantitative trait locus in yeast. Nature 416, 326–330. 10.1038/416326a.

Swinnen, E., Wanke, V., Roosen, J., Smets, B., Dubouloz, F., Pedruzzi, I., Cameroni, E., De Virgilio, C., and Winderickx, J. (2006). Rim15 and the crossroads of nutrient signalling pathways in Saccharomyces cerevisiae. Cell Div 1, 3. 10.1186/1747-1028-1-3.

Szklarczyk, D., Gable, A.L., Nastou, K.C., Lyon, D., Kirsch, R., Pyysalo, S., Doncheva, N.T., Legeay, M., Fang, T., Bork, P., et al. (2021). The STRING database in 2021: customizable protein-protein networks, and functional characterization of user-uploaded gene/measurement sets. Nucleic Acids Res 49, D605–D612. 10.1093/nar/gkaa1074.

Tanner, L.B., Goglia, A.G., Wei, M.H., Sehgal, T., Parsons, L.R., Park, J.O., White, E., Toettcher, J.E., and Rabinowitz, J.D. (2018). Four Key Steps Control Glycolytic Flux in Mammalian Cells. Cell Syst 7, 49–62 e48. 10.1016/j.cels.2018.06.003.

Teng, X., and Hardwick, J.M. (2019). Whi2: a new player in amino acid sensing. Curr Genet 65, 701–709. 10.1007/s00294-018-00929-9.

Tran, D.M., Ishiwata-Kimata, Y., Mai, T.C., Kubo, M., and Kimata, Y. (2019). The unfolded protein response alongside the diauxic shift of yeast cells and its involvement in mitochondria enlargement. Sci Rep 9, 12780. 10.1038/s41598-019-49146-5.

Traxler, L., Herdy, J.R., Stefanoni, D., Eichhorner, S., Pelucchi, S., Szucs, A., Santagostino, A., Kim, Y., Agarwal, R.K., Schlachetzki, J.C.M., et al. (2022). Warburg-like metabolic transformation underlies neuronal degeneration in sporadic Alzheimer’s disease. Cell Metab 34, 1248–1263 e1246. 10.1016/j.cmet.2022.07.014.

Vidal, M., Cusick, M.E., and Barabasi, A.L. (2011). Interactome networks and human disease. Cell 144, 986–998. 10.1016/j.cell.2011.02.016.

Wallace, E.W., Kear-Scott, J.L., Pilipenko, E.V., Schwartz, M.H., Laskowski, P.R., Rojek, A.E., Katanski, C.D., Riback, J.A., Dion, M.F., Franks, A.M., et al. (2015). Reversible, Specific, Active Aggregates of Endogenous Proteins Assemble upon Heat Stress. Cell 162, 1286–1298. 10.1016/j.cell.2015.08.041.

Weiss, C.V., Roop, J.I., Hackley, R.K., Chuong, J.N., Grigoriev, I.V., Arkin, A.P., Skerker, J.M., and Brem, R.B. (2018). Genetic dissection of interspecific differences in yeast thermotolerance. Nat Genet 50, 1501–1504. 10.1038/s41588-018-0243-4.

Workman, J.J., Chen, H., and Laribee, R.N. (2016). Saccharomyces cerevisiae TORC1 Controls Histone Acetylation by Signaling Through the Sit4/PP6 Phosphatase to Regulate Sirtuin Deacetylase Nuclear Accumulation. Genetics 203, 1733–1746. 10.1534/genetics.116.188458.

Wu, C.Y., Bird, A., Winge, D., and Eide, D. (2008). Zinc deficiency response in Saccharomyces cerevisiae: Characterization of the Zap1-responsive regulon. Faseb Journal 22. 10.1096/fasebj.22.1_supplement.782.14.

Yang, Y., Foulquie-Moreno, M.R., Clement, L., Erdei, E., Tanghe, A., Schaerlaekens, K., Dumortier, F., and Thevelein, J.M. (2013). QTL analysis of high thermotolerance with superior and downgraded parental yeast strains reveals new minor QTLs and converges on novel causative alleles involved in RNA processing. PLoS Genet 9, e1003693. 10.1371/journal.pgen.1003693.

You, C., Okano, H., Hui, S., Zhang, Z., Kim, M., Gunderson, C.W., Wang, Y.P., Lenz, P., Yan, D., and Hwa, T. (2013). Coordination of bacterial proteome with metabolism by cyclic AMP signalling. Nature 500, 301–306. 10.1038/nature12446.

Zaman, S., Lippman, S.I., Schneper, L., Slonim, N., and Broach, J.R. (2009). Glucose regulates transcription in yeast through a network of signaling pathways. Mol Syst Biol 5, 245. 10.1038/msb.2009.2.

Zelezniak, A., Vowinckel, J., Capuano, F., Messner, C.B., Demichev, V., Polowsky, N., Mulleder, M., Kamrad, S., Klaus, B., Keller, M.A., and Ralser, M. (2018). Machine Learning Predicts the Yeast Metabolome from the Quantitative Proteome of Kinase Knockouts. Cell Syst 7, 269–283 e266. 10.1016/j.cels.2018.08.001.

Zhang, A., Shen, Y., Gao, W., and Dong, J. (2011). Role of Sch9 in regulating Ras-cAMP signal pathway in Saccharomyces cerevisiae. FEBS Lett 585, 3026–3032. 10.1016/j.febslet.2011.08.023.

Zhang, J., Zhao, J., Dahan, P., Lu, V., Zhang, C., Li, H., and Teitell, M.A. (2018). Metabolism in Pluripotent Stem Cells and Early Mammalian Development. Cell Metab 27, 332–338. 10.1016/j.cmet.2018.01.008.

Zhang, N., and Cao, L. (2017). Starvation signals in yeast are integrated to coordinate metabolic reprogramming and stress response to ensure longevity. Curr Genet 63, 839–843. 10.1007/s00294-017-0697-4.

